# Arginine Di-methylation of RIPK3 Safeguards Necroptosis for Intestinal Homeostasis

**DOI:** 10.1101/2024.02.21.581356

**Authors:** Pan Zhao, Hanjun Dan, Yazhou Wang, Xin Chen, Xiangling Jiang, Yao Shen, Jiajia Wang, Zhiwei Yang, Jiasheng Zhao, Yingying Zhang, Jianyong Zheng, Wen Liu, Jian Zhang

## Abstract

The necroptosis mediated by RIPK3 is stringently regulated for intestinal homeostasis. Here we found that mice lacking *Prmt5* (Protein arginase methyltransferase 5) in intestinal epithelial cells (IECs) caused premature death with IECs necroptosis, villus atrophy and loss of Paneth cells. This pathology can be partially rescued by antibiotic treatment, germ-free breeding condition and pharmaceutical inhibition of RIPK1 and RIPK3, but aggravated for embryonic lethality by *Caspase*-8 deficiency, which demonstrating the importance of commensal bacteria and necroptosis for the *Prmt5*-IEC deficiency. Intriguingly, tumor-necrosis factor (TNF) receptor 1(*Tnfr1*) deficiency could not completely rescue the pathology, and mice deficit in Z-DNA binding protein 1(ZBP1) exhibited shorter lifespan compared with *Prmt5* null mice, suggesting *Prmt5* loss might trigger TNFR-RIPK1-depenfent and ZBP1- dependent necroptosis. Mechanically, we identified the 479-arginine residue of RIPK3 di-methylated by PRMT5 was an endogenous checkpoint for necroptosis. Furthermore, RIPK3-R479K mutation had higher affinity with both RIPK1 and ZBP1 by immunoprecipitation and STORM (Stochastic Optical Reconstruction Microscopy) analysis, which might explain the endogenous necroptosis triggered by mutated RIPK3 even without upstream stimuli. Moreover, the peptide of RIPK3-SDMA (Symmetric dimethylarginine of 479) could rescue lethality of *Prmt*5 lacking mice through necrosome formation inhibition, which demonstrating the great potential for necroptosis-related disease treatment through RIPK3 dimethylation targeting.

## Introduction

The maintenance of intestine homeostasis relies on the intestinal epithelium(IECs) which forms a physical and chemical barrier to separate microbes and toxic substances in lumen, ensuring intestinal health(Turner, 2009b). An enormous self-renewing capacity is the striking feature of the intestinal epithelium(Watson and Pritchard, 2000), with cells that have served their purpose being completely replaced by newly generated cells within 4-5 days(van der Flier and Clevers, 2009). Considering its complex structure and crucial role in maintaining intestinal homeostasis, tight control over self-renewal and cell death of the intestinal epithelium is necessary. Excessive cell death leading to barrier defects may contribute to various inflammatory diseases, including inflammatory bowel disease (IBD)(Ainsworth et al., 1989; Katz et al., 1989; Kiesslich et al., 2012; Turner, 2009a). Indeed, excessive cell death has been observed in genetic mouse models with severe chronic inflammatory pathologies as well as patients with IBD (Bosurgi et al., 2017; Farin et al., 2014; Garcia-Carbonell et al., 2018; Gunther et al., 2015; Iwamoto et al., 1996; Lehle et al., 2019; Simmons et al., 2016; Wang et al., 2020; Welz et al., 2011; Zushi et al., 1997). Therefore, understanding the mechanisms regulating IEC death is important for maintaining intestine homeostasis and elucidating the pathogenesis of gut inflammation.

Necroptosis is an alternative mode of cell death in the gut that can drive intestinal inflammation when dysregulated(Gunther et al., 2013). Inflammatory mediators such as TNFα strongly induce necroptosis signaling, suggesting a vicious cycle between epithelial cell death and dysfunctional immune response contributing to persistent intestinal inflammation seen in patients with IBD(Dechairo et al., 2001). RIP3 (also known RIPK3), a key mediator in necroptosis, belongs to the receptor-interacting protein (RIP) family of serine/threonine kinase containing RIP homotypic interaction motif (RHIM) domain(Silke et al., 2015a, b). Formation of a necrosome complex involving RIP3 and RIP1(also known as RIPK1) through their RHIM domains is necessary for inducing necroptosis in both murine and human cells (Galluzzi et al., 2017). Additionally, Z-DNA binding protein 1 (ZBP1) also known as DAI, serves as an activator of RIP3 through their RHIM domain, which senses exogenous DNA through its special domain and promotes the synthesis of type I interferon (IFN) while activating NF-κB signaling to protect cells(Kaiser et al., 2008; Zitvogel et al., 2015). The kinase activity of RIP3 is response for activating mixed lineage kinase-like (MLKL), a downstream effector protein involved in necroptosis execution(Wallach et al., 2016). Although significant progress has been made in understanding the signaling pathways of necroptosis in the gut, the intrinsic mechanism regulating RIP3 activation remains incompletely understood.

Protein arginine methyltransferase 5 (PRMT5) is a major methyltransferase that modifies symmetric di-methylation of arginine and regulates various biological processes through post-translational modification(Stopa et al., 2015). PRMT5 plays a crucial role in intestinal diseases such as enteritis and colorectal cancer development(Hernandez et al., 2023; Yang and Bedford, 2013). Mechanistic studies have suggested that PRMT5 can promote colorectal cancer cell growth by transcriptionally silencing anti-oncogenes or/and activating oncogenes(Cho et al., 2012; Prabhu et al., 2015; Shen et al., 2022; Yan et al., 2021; Zhang et al., 2015). In IBD, upregulation of PRMT5 contributes to clearing *Citrobacter rodentium* infection and promoting mucosal damage recovery during enteric infection(Hernandez et al., 2023). However, the physiological function of PRMT5 in the intestinal epithelium under homeostasis remains poorly understood.

Here, we report that mice lacking PRMT5 specifically in intestinal epithelial cells spontaneously develop severe intestinal inflammation associated with necroptosis-induced cell death leading to early mortality. This early lethality was partially rescued by inhibiting RIP1 and RIP3 activities, antibiotic treatment, maintaining germ-free conditions or deficiency in tumor-necrosis factor receptor 1(Tnfr1). This early lethality was partially rescued by inhibiting RIP1 and RIP3 activities, antibiotic treatment, maintaining germ-free conditions or deficiency in tumor-necrosis factor receptor 1(*Tnfr1*), but not CASP8 deficiency, which demonstrate indispensable role of RIP3- dependent necroptosis but not CASP8-dependent apoptosis. In addition, deficiency of *Prmt5* promoted both TNFR1-RIP1 and RIP3-ZBP1 mediated necroptosis, which was provided evidence that the mutation in arginine residues 479 of RIP3 protein di- methylated by PRMT5 significantly enhanced the interaction between RIP3 and RIP1 as well as ZBP1, respectively. Our results demonstrate that PRMT5 is crucial for the survival of intestinal epithelial cells (IECs) and maintaining gut homeostasis by protecting IECs from RIP3-mediated necroptosis.

## Results

### Epithelial PRMT5 depression triggers intestinal homeostasis breakdown

Disruption of intestinal homeostasis is likely to contribute to the development of inflammatory diseases(Garrett et al., 2010). In order to identify key molecules involved in maintaining intestinal homeostasis, we conducted a comprehensive analysis of whole genome transcriptome sequencing data obtained from patients with inflammatory bowel disease (IBD) (VanDussen et al., 2018). Our analysis revealed a significant decrease in PRMT5 expression levels in intestinal tissues from IBD patients compared to normal controls (Fig.1A). This reduction was further confirmed in biopsy specimens collected from IBD patients (Fig. 1B-C). These findings strongly suggest the involvement of PRMT5 in regulating intestinal homeostasis.

**Figure 1.**
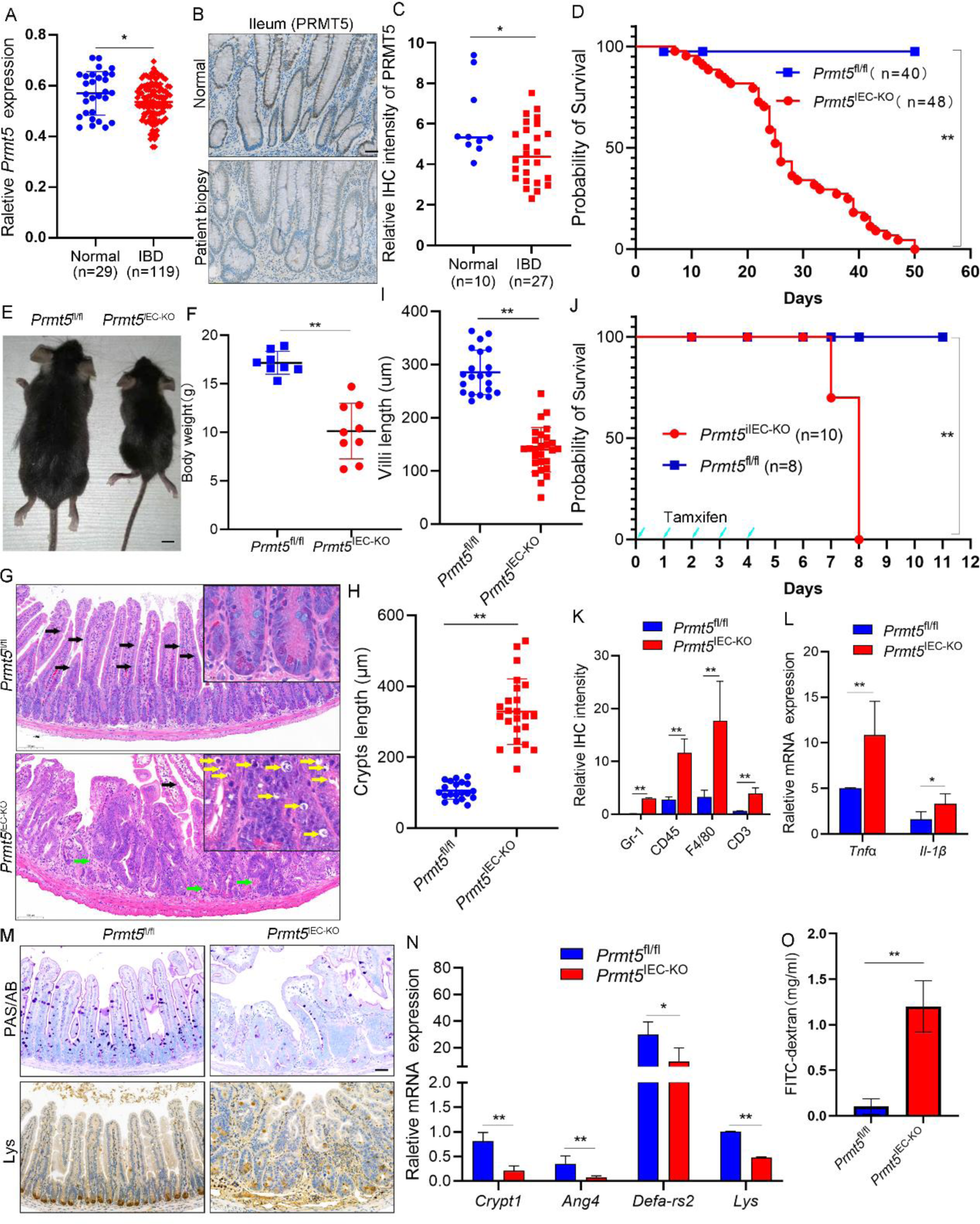
Death and spontaneous gut inflammation in mice with PRMT5 deletion in IECs A, The expression of *PRMT5* in patients with IBD during phase of inflammation (n=119) and controls (n=29), which revealed by analyzing RNA-sequencing data (Gene Expression Omnibus, GSE179285); B, The protein expression of PRMT5 was examined in normal intestinal tissues (n=10) and IBD biopsies (n=27); C, Statistical analysis was performed to compare the levels of PRMT5 protein expression; D, Cumulative survival rate analysis showed a significant difference between *Prmt5*^IEC-KO^ mice (n=48) compared to *Prmt5*^fl/fl^ mice (n=40); *P<0.05, **P<0.01. E, Macroscopic features of *Prmt5*^IEC-KO^ and *Prmt5*^fl/fl^ mice were evaluated; Scale bars, 100 µm; F, Body weight comparison between *Prmt5*^IEC-KO^ mice at 4 weeks old (n=9) and sex- matched littermate *Prmt5*^fl/fl^ mice (n=8); *P<0.05, **P<0.01. G, Haematoxylin and eosin staining (H&E staining) of small intestine sections from *Prmt5*^IEC-KO^and *Prmt5*^fl/fl^ mice. Black arrows indicate goblet; Green arrows indicate infiltrating leukocytes; Yellow arrows indicate dead cells; Scale bars, 100 µm; H, Statistical analysis for lengths of small intestine crypts; *P<0.05, **P<0.01. I, Statistical analysis for lengths of small intestine villi; *P<0.05, **P<0.01. J, Survival rates were compared among different groups: *Prmt5*^fl/fl^, *Prmt5*^iIEC-KO^ treated with five daily tamoxifen injections (75mg/kg); * P < 0 .05, ** P < 0 .01. K, Statistical analysis for Lymphocyte IHC signals in small intestine of *Prmt5*^IEC-KO^ (n=3) and *Prmt5*^fl/fl^ (n=3) mice; * P < 0 .05, ** P < 0 .01. L, Quantitative real-time PCR (qPCR) analysis of inflammatory cytokine *Tnfα* and *Il-1β* on RNA extracted from small intestine mucosa of *Prmt5*^IEC-KO^ (n=3) and control *Prmt5*^fl/fl^ (n=3) mice; *P<0.05, **P<0.01. M, Periodic acid–Schiff/Alcian blue (PAS/AB; goblet cells) staining for goblet cells, and lysozyme (Lys) immunohistochemical staining or Paneth cells in the small intestine sections. Three individual experiments were repeated; Scale bars, 100 µm; N, qPCR analysis was conducted to examine antimicrobial peptides on RNA extracted from small intestine mucosa samples both groups: *Prmt5*^IEC-KO^ (n=3) and control *Prmt5*^fl/fl^ (n=3) mice. Crypt1, cryptidin 1; LysP, lysozyme P. *P<0.05, **P<0.01. O, Intestinal permeability assay in *Prmt5*^IEC-KO^ (n=3) and *Prmt5*^fl/fl^ (n=3) mice using FITC-labelled dextran. *P<0.05, **P<0.01.

The role of PRMT5 in regulating intestinal homeostasis has not been fully understood yet. Therefore, we generated a mouse line with conditional knockout of PRMT5 (*Prmt5*^fl/fl^) using loxP technology (Extended Data Fig. 1A). Specifically deletion *Prmt5* in the mice intestinal epithelium by generating VIL-Cre*Prmt5*^fl/fl^ (*Prmt5*^IEC-KO^), which progeny were born at abnormal Mendelian ratio (Extended Data Fig. 1B). Protein analysis of various organs form *Prmt5*^IEC-KO^ mice confirmed completely *Prmt5* deletion in small intestine and partially deletion in colon (Extended Data Fig. 1C). Approximately 70% of *Prmt5*^IEC-KO^ mice died within 50 days (Fig. 1D) and exhibited reduced body weight (Fig. 1E-F and Extended Data Fig. 3A) along with severe enteritis symptoms (Extended Data Fig. 1D-E).

The small intestine of surviving *Prmt5*^IEC-KO^ mice exhibits abnormal tissue architecture and aberrant crypts (Fig.1H-J). However, the development of the gut in *Prmt5*^IEC-KO^ mouse embryos did not display obvious abnormalities (Extended Data Fig. 3B). When VIL-CreERT2*Prmt5*^fl/fl^ (*Prmt5*^iIEC-KO^) mice were utilized to completely delete *Prmt5* in the intestinal epithelium after tamoxifen treatment (Extended Data Fig. 1H), all the mice dead within 3 days after tamoxifen injection was done (fig.1J) and also displays abnormal tissue architecture and aberrant crypts (Extended Data Fig.1J- L). Thus, the phenotype of *Prmt5*^IEC-KO^ mice is not specific to neonate. The small intestine of *Prmt5*^IEC-KO^ exhibited inflammation characterized by the presence of inflammatory infiltrates (Fig 1K and Extended Data Fig. 1I), increased levels of inflammatory factors *TNFα*, *Il-1β* (Fig.1L), loss of mucus-producing goblet cells and antimicrobial peptide (AMP)-producing Paneth cells (Fig. 1M). Accordingly, expression of the antimicrobial peptides and cryptidin 1 were decreased (Fig.1N), which combined with impaired antimicrobial ability resulted in physical barrier dysfunction and intestinal leakage, as shown by decreased tight junctions (Extended Data Fig. 1G) and increased fluorescence (Fig.1O). Our data suggested deletion PRMT5 in intestine results in homeostasis breakdown.

### PRMT5 inhibits necroptosis of intestinal epithelial cells

Massive intestinal cell death occurs after *Prmt5* deletion, as demonstrated shown by H & E staining and terminal deoxynucleotidyl transferase dUTP nick end labelling (TUNEL) (Fig. 1G and Extended Data Fig.2A). To explore forms of cell death among IECs when there is a deficiency in *Prmt5*, we observed massive necroptotic cells within subcellular features using transmission electron microscopy (Fig.2A-B). The increased phosphorylated levels RIP1, phosphorylated RIP3 and phosphorylated MLKL-the hallmarks of necroptosis-were detected within intestines from *Prmt5* deficiency mice (Fig.2C-D and Extended Data Fig.2C).

**Figure 2.**
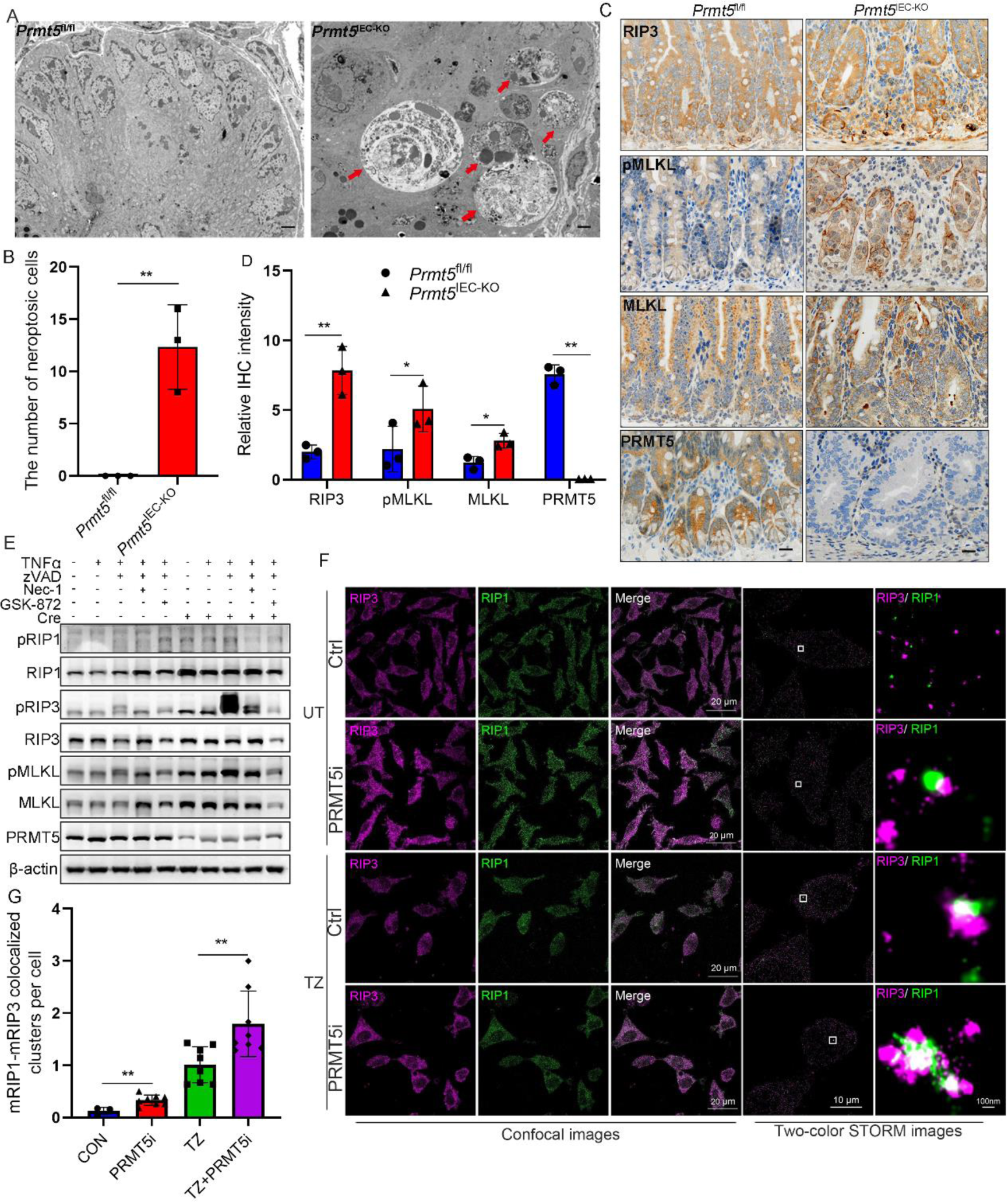
Epithelial PRMT5 depression triggers necroptosis A, Electron microscopy images of necrotic-like IECs in *Prmt5*^IEC-KO^ (n=3) mice. Red arrows: necrotic-like IECs. TEM, transmission electron microscopy; B, Statistical analysis for necrotic-like IECs in *Prmt5*^IEC-KO^ mice; *P<0.05, **P<0.01. C, Necroptosis IHC signals of phosphorylation level of MLKKL and total RIP3 in small intestine of *Prmt5*^IEC-KO^ (n=3) and *Prmt5*^fl/fl^ (n=3) mice; Scale bars, 100 μm; D, Statistical analysis for necroptosis IHC signals in *Prmt5*^IEC-KO^ mice. *P<0.05, **P<0.01. E, Immunoblotting of phosphorylation level of RIP1, RIP3 and MLKL in primary mouse embryonic fibroblasts (MEFs) derived from *Prmt5*^fl/fl^ mice; F, V5-RIP3 or Flag-RIP1 were overexpressed in 293T cells and treated as indicated (DMSO; 0.5 μM PRMT5i; T: 20 ng^-ml^ TNFα; Z: 20 μM zVAD). Confocal and two- colour STORM images of RIP1–RIP3 complex in cells; G, Statistical analysis for RIP3-RIP1colocalized clusters per cell in two-colour STORM imaging. *P<0.05, **P<0.01.

We investigated the role of *Prmt5-*mediated necroptosis in cellular systems. The conditional deletion of *Prmt5* using CRE-inducible mouse embryonic fibroblasts (MEFs) resulted in necroptosis both in the presence and absence necroptosis induced by TZ (TNFα combined with zVAD-FMK,a pan-caspase inhibitor). This effect was rescued by the RIP1-specific inhibitor necrostatin-1s (Nec-1s) or the RIP3-specific inhibitor GSK-872, as indicted by western blotting and TUNEL staining (Fig.2E and Extended Data Fig.2D-E). To further assess the PRMT5 kinase-specific inhibitor JNJ-64619178 (PRMT5i), we employed L929 mouse fibroblast cell line to trigger necroptosis. The western data showed that PRMT5i could accelerate necroptosis signaling in the presence of TZ, which was confirmed by transmission electron microscopy and TUNEL staining (Extended Data Fig.2G-I).

The initiation of induction of necrosis through RIP3 activation involved a heterodimeric filamentous structure known as a necrosome, mediated by assembly between RIP3 and RIP1 via their respective RHIM domains(Li et al., 2012). To investigate the role of PRMT5 in cellular necrosome, we imaged RIP1-RIP3 complexes in L929 cells using confocal microscopy and two-color STORM imaging, which revealed that PRMT5i increased both the numbers and size of necrosomes regardless of whether or not they were induced for necroptosis (Fig2F-G). These results also supported by co-immunoprecipitation experiments showing interaction between RIP1 and RIP3 proteins (Extended Data Fig.2J). In conclusion, mice lacking PRMT5 expression specifically in IECs spontaneously developed severe intestinal inflammation associated with IECs undergoing programmed cell death via necroptosis leading to early mortality.

### Necroptosis responses contribute pathologically to *prmt5* deficiency but can occur independently from TNFR1-RIP1 signaling

To assess the pathological contribution of necroptosis, we employed GKS-872 as a RIP3 inhibitor (since *Prmt5* and *Rip3* genes are located on the same chromosome) to effectively block necroptosis in vivo. Intraperitoneal injection of GSK-872 significantly prolonged life time and intestine length in *Prmt5*^iIEC-KO^ mice (Fig.3A-B). Histological analysis of ileal sections from GSK-872 treated mice revealed that partial restoration of tissue architecture, overall reappearance of Paneth cells, and reduced infiltration of CD45+ immune cells (Fig.3C-D). Additionally, it has been reported that mice with IEC-specific caspase-8 deficiency (*Casp8*^IEC-KO^) develop ileitis characterized by loss of Paneth cells due to necroptosis(Gunther et al., 2011). Interestingly, *Prmt5*^iIEC-KO^ *Casp8*^iIEC-KO^ DKO (double knockout) mice exhibited shorter lifespans compared to *Prmt5*^iIEC-KO^ mice (Fig.3A). Histological examination showed no significant difference in tissue architecture and number of Paneth cells (Lys) between *Prmt5*^iIEC-KO^ *Casp8*^iIEC-KO^ DKO mice and *Prmt5*^iIEC-KO^ mice; however, there was a marked increase in CD45+ immune cell infiltrates and phosphorylated MLKL immunohistochemistry intensity in the former group (Fig.3E-F). Collectively, these findings suggested that necroptosis may contribute to the pathological phenotype observed in *Prmt5* deficiency mice.

**Figure 3.**
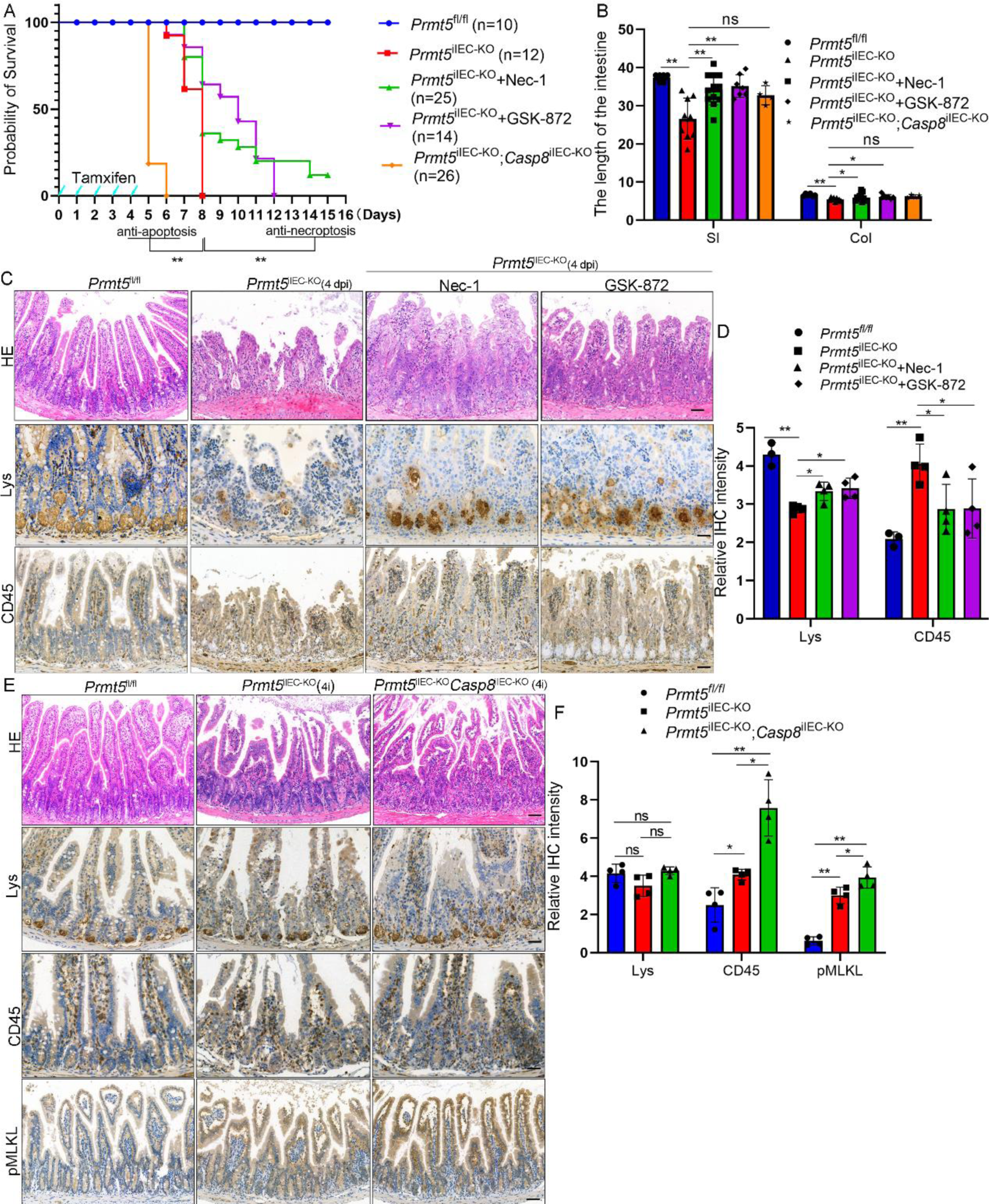
Lethality of *Prmt5*^iIEC-KO^ in mice rescued by inhibition of RIP1 and RIP3, but not *Casp8*-difciency A, Survival rates were compared among different groups: *Prmt5*^iIEC-KO^ mice treated with Nec-1 or GSK-872, *Prmt5*^iIEC-KO^*Casp8*^iIEC-KO^, and *Prmt5* ^fl/fl^ mice. **P<0.01. B, Statistical analysis for the gut length in different groups of mice. SI (small intestine), Col (colon); *P<0.05, **P<0.01 C, Representative images of small intestine sections from mice with the indicated genotypes stained with HE or immunostained for Lys and CD45. Scale bars, 100 μm. D, Statistical analysis for Lys and CD45 IHC signals; *P<0.05, **P<0.01 E, Representative images of small intestine sections from mice with the indicated genotypes stained with HE or immunostained for Lys, CD45 and pMLKL. Scale bars, 100 μm; F, Statistical analysis for Lys, CD45 and pMLKL IHC signals; *P<0.05, **P<0.01

Bacterial colonization shortly after birth is tightly associated with cell death, gut inflammation, and mortality(Takahashi et al., 2014). Additionally, RIP1 is required for the induction of a prototypical model of necroptosis by the proinflammatory cytokine TNF(Pasparakis and Vandenabeele, 2015). We hypothesized *Prmt5* deletion sensitizes IECs to TNF, which responds to commensal bacteria and triggers necroptosis. To this hypothesis, we used a cocktail of broad-spectrum antibiotics (ABX) and germ-free mice to clear away commensal bacteria or mice deficient in TNFR1(*Tnfr*^-/-^) to block upstream of necroptosis. Unexpectedly, deficiency in TNFR1 did not increase body weight compared with *Prmt5* deletion (Extended Data Fig.4A). Bloking commensal bacteria or TNF signaling did not completely rescued the mortality and the length of intestine in *Prmt5*^iIEC-KO^ mice (Extended Data Fig.4B-C). Histological examination revealed that blocking commensal bacteria and inflammation strongly ameliorated-but did not fully prevent-the pathological phenotype as shown by partial re-appearance of Paneth cells, decreased RIP3 immunohistochemistry intensity, reduced CD45+ immune cell infiltrates but tissue architecture was not restored (Extended Data Fig.4D-E).Similar to epithelial TNFR1 deficiency, inhibition of RIP1 kinase activity by Nec-1 also strongly increased lifespan and intestine length in *Prmt5*^iIEC-KO^ mice (Fig.3A-B). Histological examination revealed partial restored tissue architecture, overall re-appearance of Paneth cells and reduced CD45+ immune cell infiltrates (Fig. 3C-D). Collectively, these results revealed that while TNFR1-RIP1 signaling plays an important role in pathogenesis in *Prmt5*^iIEC-KO^ mice; there are also other mechanisms independent from this pathway contributing to it.

### PRMT5 suppress necroptosis by catalyzing the di-methylation of RIP3

The machines through which PRMT5 suppresses necroptosis were investigated by identifying potential di-methylation proteins in the intestine of *Prmt5*^IEC-KO^ mice using mass spectrometry analysis. Kyoto Encyclopedia of Genes and Genomes (KEGG) analysis was employed to comprehensively describe the changes in signaling pathways, revealing a significant increase in necroptosis signaling in *Prmt5*^IEC-KO^ mice (Fig.4A). Additionally, two arginine residues (R477 and R479) on RIP3 were identified as being di-methylated with high confidence (Fig.4B), which was confirmed by immunoprecipitation with an antibody against RIP3 followed by immunoblotting with an antibody against Symmetric Di-Methyl Arginine Motif (SDMA) using mutant E444Q to inhibit PRMT5 enzymatic activity (Fig 4C). Furthermore, Furthermore, we demonstrated that R479 of RIP3 is specifically methylated by PRMT5 (Fig.4D). Notably, R479 is highly conserved among different isoforms of RIP3 proteins, suggesting its methylation may be functionally important. In this study, we focused on investigating the biological function of di-methylation at R479 (R479me2) in RIP3 protein. A specific antibody targeting R479me2 (RIP3^SDMA479^) was generated for rapid and precise detection of methylation at R479 of RIP3 protein. This antibody was used to demonstrate that treatment with a PRMT5 inhibitor abolished methylation at R479 of RIP3 protein via IP assay (Fig.4E). Immunoblotting and histologic immunohistochemistry examinations further confirmed that *Prmt5* deficiency significantly reduced methylation at R479 of RIP3 protein to a greater extent (Extended Data Fig.5A-C).

**Figure 4.**
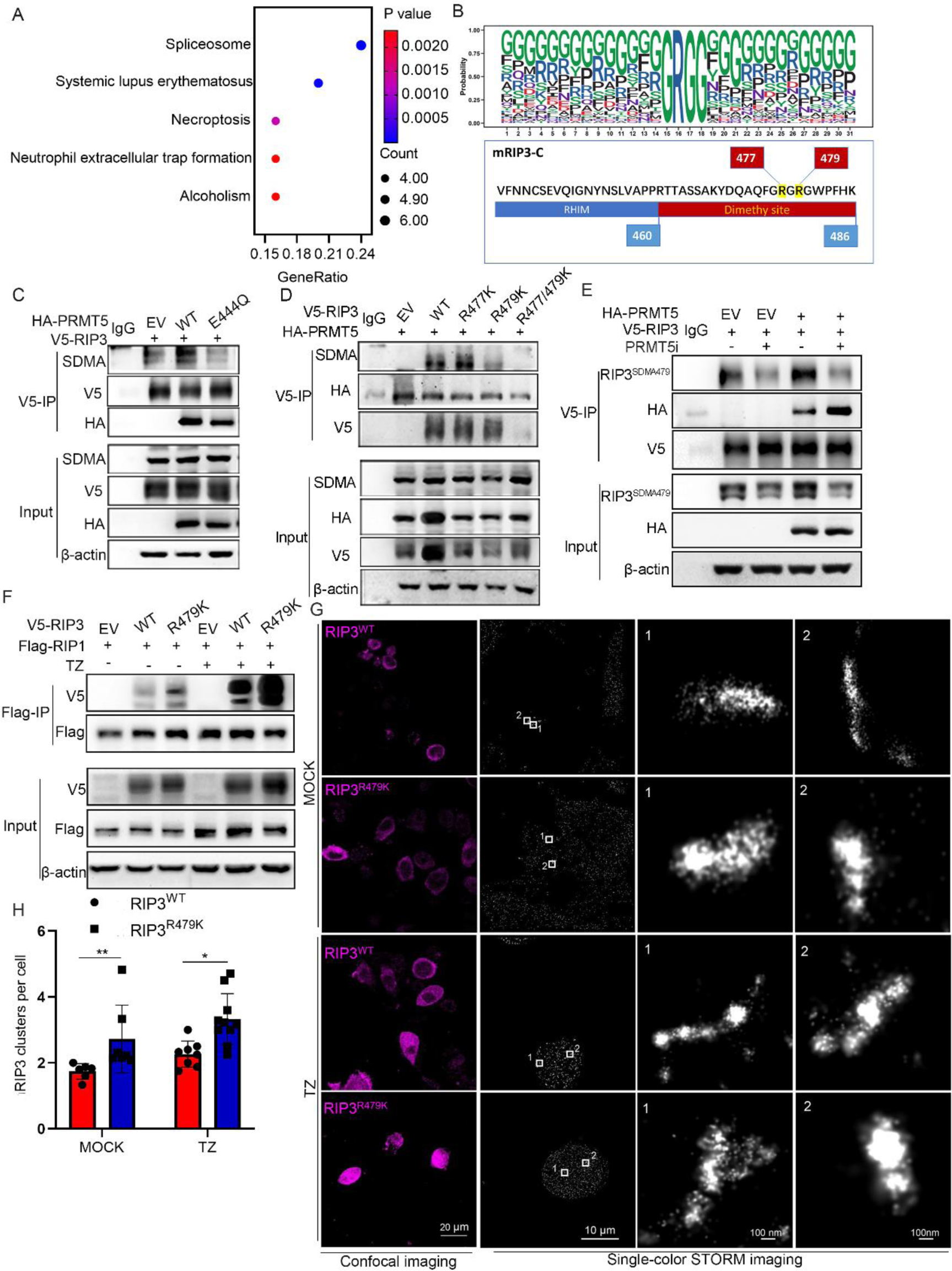
Arginine methylation of RIP3 on R479 suppresses necroptosis mediated by RIP1 A, The arginine methylation identified through mass spectrometry analysis of IECs isolated from *Prmt5* ^fl/fl^ or *Prmt5*^IEC-KO^ mice was subjected to KEGG analysis(n=6); B, Two potential Arginine methylation sites at R477 and R479 in C-terminal fragment of RIP3; C, V5-RIP3 together with HA-PRMT5 WT or enzyme-dead E444Q mutant overexpressed in 293T cells and immunoprecipitated with anti-V5 beads, followed by immunoblotting with anti-symmetric di-methyl arginine motif (SDMA) antibody; D, HA-PRMT5 together with V5-RIP3 WT or arginine methylation of RIP3 on R477, R479 or double mutant overexpressed in 293T cells, immunoprecipitated with anti- V5 beads, followed by immunoblotting with anti-SDMA antibody; E, V5-RIP3 together with HA-PRMT5 overexpressed in 293T cells treated with or without PRMT5i and immunoprecipitated with anti-V5 beads, followed by immunoblotting with anti-SDMA of RIP3 on R479 (RIP3^SDMA479^) antibody; F, Flag-RIP1 together with V5-RIP3 WT or arginine methylation of RIP3 on R479 mutant R479K overexpressed in 293T cells with or without necroptosis induction (T: 20 ng^-ml^ TNFα; Z: 20 μM zVAD); G, V5-RIP3 WT or mutant R479K were expressed in RIP3-KO L929 cells and treated as indicated (DMSO, 20 ng^-ml^ TNFα, 20 μM zVAD). Confocal and signal-colour STORM images showed the RIP3 necrosome during cell necroptosis. The white box magnified images highlighting the RIP3 necrosome; H, Statistical analysis for RIP3 clusters per cell; *P<0.05, **P<0.01

To elucidate the biological function of methylation on R479 in RIP3 protein, a mutant form of RIP3 carrying an amino acid substitution at position 479(RIP3^R47K^) was utilized for functional exploration purposes. The introduction of this mutation not only scientifically increased the present of cell death in RIP3-KO L929 cells treated with or without TZ (Extended Data Fig.5D), but also enhanced necroptosis signaling compared with RIP3^WT^ in RIP3-KO L929 cells (Extended Data Fig.5E). The combination RIP3 with RIP1 was initiation of induction of necrosis. Inhibition of PRMT5 promotes the interaction between RIP3 and RIP1 (Extended Data Fig.5F). Consistently, IP analysis in HEK293T cells and RIP3-KO L929 cells showed that RIP3^R479K^ enhances the ability of RIP3 to interact with RIP1 (Fig.4E and Extended Data Fig.5G-H). Additionally, we performed MD simulations to model the structures of both wild-type (WT) and R479K mutant forms of RIP3 in complex with RIP1. The binding energies (*ΔGbind*) and backbone-atom root-mean-square deviations (RMSD) indicated that RIP3^R479K^ has a higher propensity to form a necrosome complex with RIP1 compared to RIP3^WT^ (Extended Data Fig.5J-K). Furthermore, our modeling results demonstrated that the interaction between RIP3^R479K^ and RIP1 is less constrained than that between RIP3^WT^ and RIP1 (Extended Data Fig.6A-C). We also analyzed various structural parameters such as radius of gyration (Rg), hydrogen bond (H-bond), solvent accessible surface area (SASA), and contact number over time, which collectively supported the stronger binding ability of RIP3^R479K^ towards RIP1 compared to RIP3^WT^ (Extended Data Fig.6D-G). Moreover, confocal microscopy images combined with single-color STORM imaging revealed that overexpression of RIP3^R479K^ in RIP3-knockout L929 cells resulted in increased oligomeric assembly of RIP3 compared to RIP3^WT^ (Fig.4G- H), which is essential for triggering substantial cell death exhibiting necroptotic characteristics (Chen et al., 2022a). Therefore, *Prmt5* deficiency leads to cell death through activation dependent on RIP3.

### PRMT5 suppress necroptosis mediated by RIP3-ZBP1

We have demonstrated that there is a TNFR1-RIP1-independent mechanism contribute to the pathogenesis in *Prmt5*^iIEC-KO^ mice. ZBP1 also acts as a key mediator, interacting with RIP3 through its RHIM domain to trigger necroptosis(Lin et al., 2016; Newton et al., 2016). We hypothesized that a ZBP1-RIP3-dependent mechanism might contribute to the pathogenesis in *Prmt5*^iIEC-KO^ mice. KEGG analysis was used to fully describe the intracellular environment required for activating ZBP1-mediated necroptosis, revealing the simultaneous activation of multiple viral infections and antiviral host defense pathways (Fig.5A). Inhibition PRMT5 promoted the interaction between ZBP1 and RIP3 in HEK293T cells (Fig.5B). The RIP3^R479K^ mutant exhibited increased interaction with ZBP1 both in HEK293T cells and RIP3-KO L929 cells (Fig.5C-D). Two-colour confocal and STROM images showed more clusters indicating colocalization of RIP3^R479K^ and ZBP1 compared to RIP3^WT^ in RIP3-KO L929 cells (Fig.5E-F and Extended Data Fig.7C-D). A mutated RHIM domain in ZBP1 protein(192-195IQIG- AAAA) was used as negative control, which revealed that colocalization of RIP3^R479K^ and ZBP1-ΔRHIM1 colocalization occurred more frequently than RIP3^WT^ in RIP3-KO L929 cells (Extended Data Fig.7A-D). Additionally, we performed structural modeling of both WT/R479K forms of RIP3 and ZBP1 using MD simulation; binding energies (ΔGbind) and backbone-atom root-mean-square deviations (RMSD) suggested it was easier for RIP3^R479K^ to combine with ZPB1 forming necrosome complexes (Extended Data Fig.7E-G). Collectively, these data suggest ZBP1-RIP3 signaling was indispensable in pathogenesis caused by *Prmt5* deletion. Unexpectedly, genetic double ablation of both Zbp*1* and *Prmt5*(*Prmt5*^iIEC-KO^ *Zbp1*^-/-^) in IECs resulted in shorter lifespans than *Prmt5*^iIEC-KO^ (Fig.5G). The ZBP1 protein can activate PANoptosis, which is regulated by a complex with key features of apoptosis, pyroptosis, and necroptosis(Karki et al., 2021). Therefore, we investigated the signaling pathways involved in apoptosis and pyroptosis and discovered that pyroptosis was induced in ZBP1-KO L929 cells by RIP3^R479K^ and PRMT5i (Extended Data Fig.8A-B). Disulfiram, a specific inhibitor for pyroptosis, significantly prolonged the lifespan of *Prmt5*^iIEC-KO^ *Zbp1*^-/-^ mice (Fig.5G). These findings suggest that necroptosis mediated by ZBP1-RIP3 signaling plays an essential role in the pathogenesis in *Prmt5* deletion; however, genetic double ablation of *Zbp1* and *Prmt5* triggers both necroptosis and pyroptosis.

**Figure 5.**
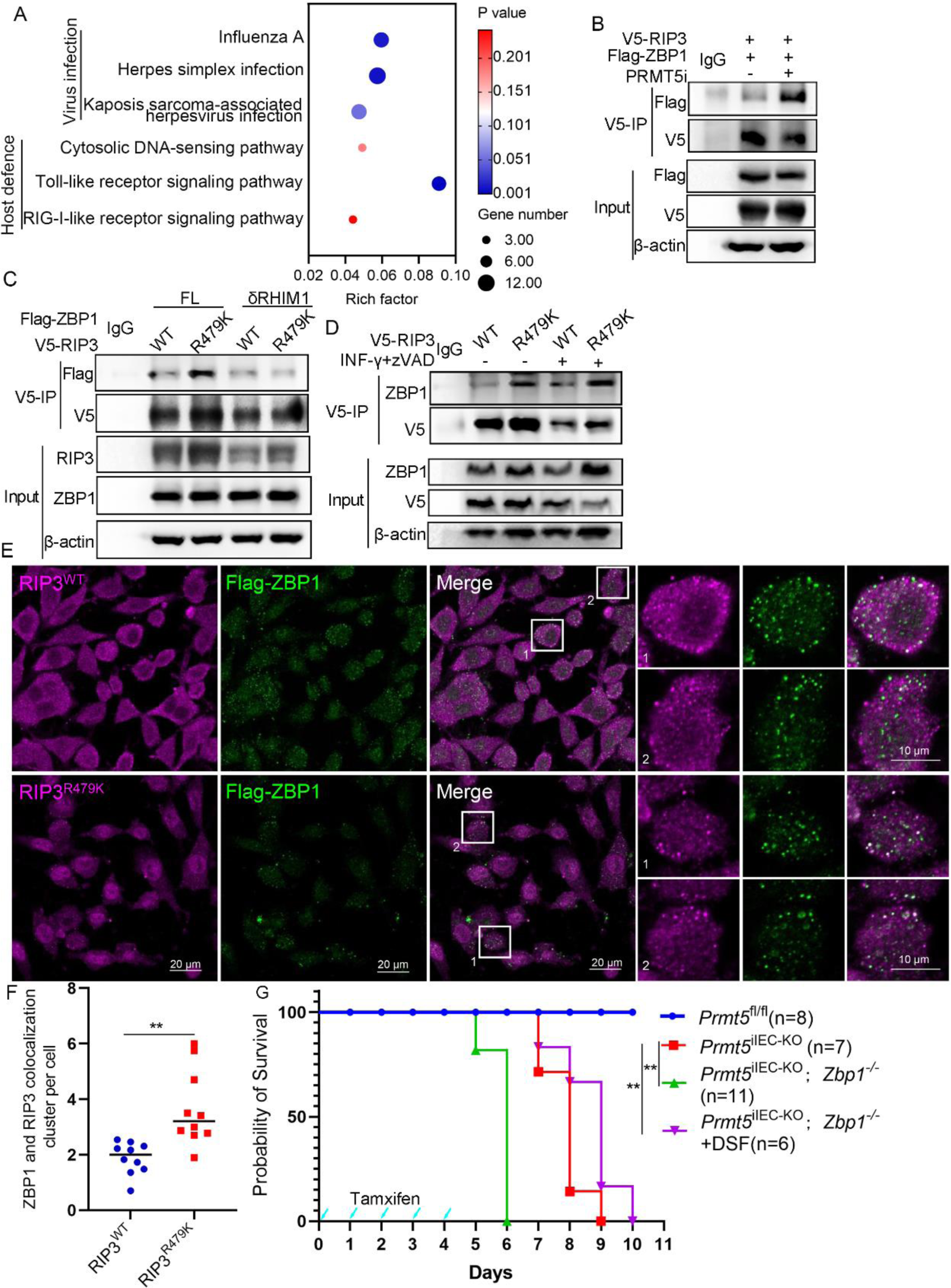
Arginine methylation of RIP3 on R479 suppresses necroptosis mediated by ZBP1. A, KEGG analysis for single cell transcriptome of IECs derived from *Prmt5* ^fl/fl^ or *Prmt5*^IEC-KO^ mice (n=2) showed upregulated differentially expressed genes associated with multiple viral infection and host defence responses; B, V5-RIP3 together with Flag-ZBP1 were overexpressed in 293T cells and treated as 0.5 μM PRMT5i and immunoprecipitated with anti-V5 beads, followed by immunoblotting with Flag antibody; C, Flag-ZBP1 WT or RHIM domain mutant (δRHIM1) together with V5-RIP3 WT or mutant R479K overexpressed in 293T cells and immunoprecipitated with anti-V5 beads, followed by immunoblotting with Flag antibody; D, V5-RIP3 were overexpressed in RIP3-KO L929 cells and treated as indicated (50 ng^-ml^ IFNγ, 20 μM zVAD), immunoprecipitated with anti-V5 beads, followed by immunoblotting with ZBP1 antibody; E, Two-colour confocal images of RIP3 WT or mutant R479K and ZBP1 in cells, the white box magnified images highlighting the RIP3 and ZBP1 colocalization; F, Statistics analysis for RIP3-ZBP1 colocalized clusters per cell in two-colour imaging. *P<0.05, **P<0.01. G, Survival rates were compared among different groups: *Prmt5*^fl/fl^, *Prmt5*^iIEC-KO^*zbp1*^iIEC-KO^ and *Prmt5*^iIEC-KO^*zbp1*^iIEC-KO^ treated with DSF;

### Di-methylation of RIP3 in polypeptide suppress necroptosis

To test the impact of RIP3 di-methylation, we administered RIP3 polypeptide with (TAT-RIP3^SDMA^) or without demethylation (TAT-RIP3^WT^) to *Prmt5*^iIEC-KO^ mice. Treatment with TAT-RIP3^SDMA^ significantly reduced cell death and necroptosis signaling in L929 cells (Fig.6A and Extended Data Fig.9C). Colocalization of RIP3 and RIP1 clusters was significantly decreased in L929 cells treated with TAT-RIP3^SDMA^ (Fig.6B-C). Additionally, it was observed that TAT-RIP3^SDMA^ could directly decrease the binding of RIP3 to RIP1 (Fig.6D and Extended Data Fig.9D). Furthermore, treatment with TAT-RIP3^SDMA^ significantly improved overall lifetime and length of intestine of *Prmt5*^iIEC-KO^ mice (Fig.6E and Extended Data Fig.9E). Histologic examination showed that treatment with TAT-RIP3^SDMA^ greatly alleviated abnormal tissue architecture and inflammation, while also recovering the number of Paneth cells (Fig.6F-G). These results indicate that di-methylation of RIP3 strongly reduces necroptosis in *Prmt5*^iIEC-KO^ mice.

**Figure 6.**
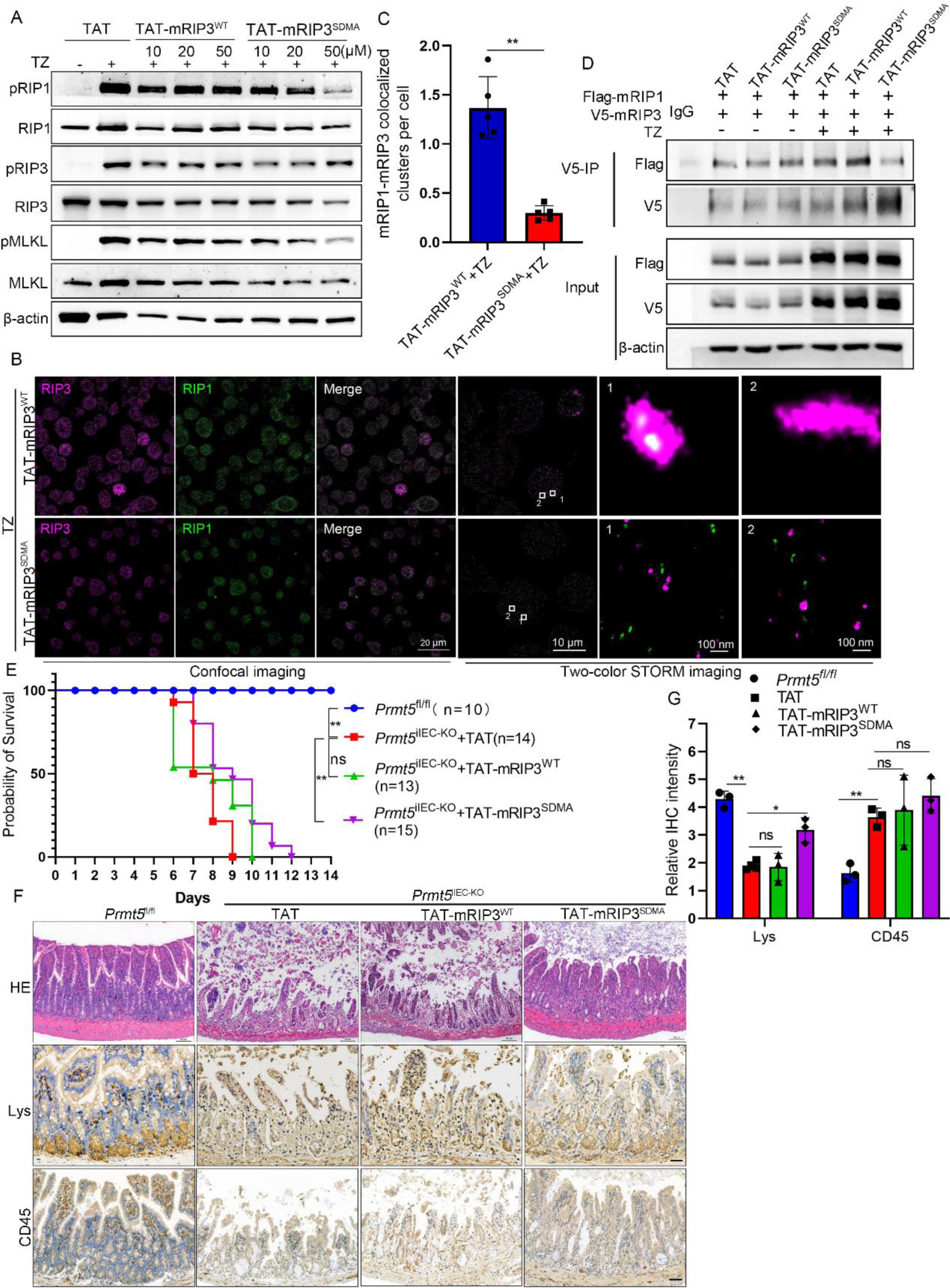
Peptide modified with arginine methylation of RIP3 on R479 suppresses necroptosis mediated by RIP3 A, Immunoblotting of phosphorylation of necroptosis signaling molecules in L929 cells and treated as indicated (50 μM peptides, 20 ng^-ml^ TNFα, 20 μM zVAD); B, Confocal and two-colour STORM images of RIP1–RIP3 complex in L929 cells that treated with TAT-RIP3 peptides with (TAT-mRIP3^WT^) or without modifying with arginine methylation at R479 on RIP3 (TAT-mRIP3^SDMA^) when necroptosis was induced by TZ (T: 20 ng^-ml^ TNFα; Z: 20 μM zVAD); C, Statistical analysis for RIP3-RIP1colocalized clusters per cell in two-colour STROM images. *P<0.05, **P<0.01. D, Both V5-RIP3 and Flag-RIP1 were overexpressed in 293T cells treated with TAT, TAT-mRIP3^WT^ or TAT-mRIP3^SDMA^ peptides with or without necroptosis induced by TZ. Immunoprecipitated with anti-V5 beads; E, Survival rate of *Prmt5*^iIEC-KO^mice treated with TAT, TAT-mRIP3^WT^ or TAT- mRIP3^SDMA^ peptides (2 mg^-kg^); *P<0.05, **P<0.01. F, Representative images of small intestine sections from mice with the indicated peptides stained with HE or immunostained for Lys and CD45; Scale bars, 100 μm. G, Statistical analysis for IHC signals of Lys and CD45; *P<0.05, **P<0.01.

## Discussion

Earlier studies have suggested that mice with insufficient levels of *Prmt5* exhibit a diminished capacity to clear intestinal infections and maintain intestinal homeostasis, which is dependent on the regulation of specific canonical crypt goblet cell gene programs(Hernandez et al., 2023). Indeed, our study has found that found a decrease in PRMT5 expression in patients with IBD, and mice lacking PRMT5 in IECs spontaneously develop severe intestinal inflammation. A previous study reported RIP3 as a target protein of PRMT5 through mass spectrometry screening of cellular proteins (Chauhan et al., 2023), but there was no evidence to prove its existence in vivo or clarify the precise mechanism by which di-methylation of RIP3 activates necroptosis. In this study, we demonstrated that di-methylation-mediated activation of IECs necroptosis by RIP3 led to early death in *Prmt5-* deficient mice. Furthermore, we proposed that loss of symmetric di-methylation arginine modification in the RIP3 protein promotes TNFR1- RIP1-dependent and ZBP1-dependent necroptosis.

The aforementioned studies raise an important question regarding the activation of necroptosis under RIP3 demethylation conditions. The formation of the necrosome, which involves RIP3 and RIP1 binding through their RHIM domain, serves as an initiator for activating necroptosis(Li et al., 2012). Interestingly, both the RHIM domain and di-methylation are present in the C-terminal region of the RIP3 protein, suggesting that PRMT5-mediated di-methylation of RIP3 may impact necrosome formation. Recent studied have reported that O-GlcNAcylation of RIP3 prevents its interaction with RIP1 via the RHIM motif(Li et al., 2019a). In our study, we found that demethylation of R479 on RIP3 enhances necrosome formation using immunoprecipitation (IP) and single- or two-STROM techniques. Unexpectedly, genetic deletion of *Tnfr1* and inhibition of RIP1 did not completely rescue early lethality in *Prmt5*- deficient mice, indicting the presence of TNFR1-RIP1 independent mechanisms. To explore this further, we investigated ZBP1 since it has been shown that RIP3 can directly bind to ZBP1 through the RHIM motif to activate necroptosis(Kaiser et al., 2008; Zitvogel et al., 2015). Indeed, demethylation at R479 on RIP3 enhanced its interaction with ZBP1; however, mice with genetic deletions in both *Zbp1* and *Prmt5* (*Prmt5*^iIEC-KO^*Zbp1^-/-^*) had a shorter lifespan compared to *Prmt5*^iIEC-KO^ mice. Additionally, inhibition of pyroptosis by DSF prolonged the lifespan of *Prmt5*^iIEC-KO^*Zbp1^-/-^* mice. These findings support ZBP1 as an activator involved in PANoptosis regulation—a complex process encompassing apoptosis, pyroptosis, and necroptosis (Karki et al., 2021). The genetic deletion models involving *Gasdermind, Zbp1* and *Prmt5* in mice are currently being studied. These findings highlight a crucial role for di-methylation modification-mediated assembly between RIP3, RIP1 and ZBP1 in activating necroptosis.

The MLKL protein, known for its role as an executor of necroptosis, plays a crucial role in both TNFR-RIP1-dependent and ZBP1-dependent necroptosis [(Wallach et al., 2016). However, genetic deletion of *Mlkl* and *Prmt5*(*Prmt5*^iIEC-KO^ *Mlkl* ^iIEC-KO^) did not rescue the morality but instead shortened the lifespan of *Prmt5*^iIEC-KO^ mice (data not shown). This raises the question of whether MLKL has a necroptosis-independent function under conditions of *Prmt5* deficiency. This question remains an area of ongoing research.

Taken together, our findings demonstrate that RIP3 plays a crucial role in the premature mortality of mice with intestinal inflammation by inducing necroptosis of PRMT5-deficient IECs. This underlying mechanism may have implications for the pathogenesis of intestinal inflammation in the context of other diseases associated with prostate through tadalafil-mediated inhibition of PRMT5 activity and cancers characterized by up-regulated PRMT5 expression(Wu et al., 2022). Additionally, our results suggest that ZBP1 could contribute to intestinal inflammation in PRMT5- deficient mice. Furthermore, we propose that di-methylation of RIP3 might serve as an intrinsic protective mechanism for IECs within an organism.

## Methods

### Mice

VIL-Cre and VIL-CreERT2 mice were purchased from Jackson Laboratory. *Tnfr1*^-/-^ mice on a C57BL/6J background were a gift from Professor Xin Lin at Tsinghua University(Noah et al., 2011). The *Zbp1*^−/−^ mice were a gift from Professor Jiahuai Han and Wei Mo at Xiamen University(Wang et al., 2020). *Casp8*^fl/fl^ and *Mlkl*^fl/fl^ mice were kind gifts from Professor Yazhou Wang at The Fourth Military Medical University(Chen et al., 2014; Gunther et al., 2011). *Prmt5*^fl/fl^ mice were generated in this study (Extended Data Fig. 1A). VIL-Cre or VIL-CreERT2 mice were crossed with *Prmt5*^fl/fl^ ^mice^ to ablate *Prmt5* in IECs. VIL-CreERT2 *Prmt5*^fl/fl^ mice were crossed with *Casp8*^fl/fl^, *Tnfr1*^−/−^, *Mlkl*^fl/fl^ or *Zbp1*^−/−^ mice to generate inducible double-knockout mice. To induce VIL–CreERT2-mediated recombination, eight-week-old mice received a 75 mg/kg dose of tamoxifen by intraperitoneal injection for 5 consecutive days. The sample size of mice for experiments was empirically determined, and the control or experimental groups were randomly assigned. Sex preference did not exist in the experiment. All mice were on a C57BL/6 background and housed in a specific pathogen-free environment at The Fourth Military Medical University Laboratory Animal Center. Sample size estimation was not performed by any statistical methods. There was no randomization, but stratification was used to achieve a similar sex ratio among the experimental groups. No blinding was performed.

### Cell culture

L929 fibroblasts were purchased from Procell Life Science & Technology Co., Ltd. (CL-0137), and *RIP3*^−/−^, *Zbp1*^−/−^, and *Mlkl*^−/−^ L929 cells were obtained from Professor Jiahuai Han’s laboratory at Xiamen University and genetically modified by the CRISPR/Cas9 or TALEN gene editing technique. The genetic modifications were confirmed by immunoblotting for the expression of the indicated proteins. The targeting sequences were as follows:

RIP3―5′-CTAACATTCTGCTGGA-3′; ZBP1―5′-AAGATCTACCACTCACGTC-3′. MLKL―5′-ATCATTGGAATACCGT-3′;

These cell lines were all cultured in Dulbecco’s modified Eagle’s medium (DMEM) with 10% (v/v) heat-inactivated fetal bovine serum (FBS), 2 mM glutamine and antibiotics (10,000 units penicillin and 10 mg streptomycin per ml) at 37°C in a humidified incubator containing 5% CO2.

### Mouse embryonic fibroblasts

Homologous *Prmt5*^fl/fl^ embryos on day 13.5 mice were used to isolate MEFs and cultured in Dulbecco’s modified Eagle’s medium (DMEM) with 10% (v/v) heat-inactivated fetal bovine serum (FBS), 2 mM glutamine and antibiotics (10,000 units penicillin and 10 mg streptomycin per ml) at 37°C in a humidified incubator containing 5% CO2.

### Western blot

Protein lysates were collected from L929 cells, MEF cells and tissue lysates. separated by SDS–PAGE and then analysed by immunoblotting. PRMT5 was detected by mouse monoclonal antibody (#79998, Cell Signaling Technology). The necroptosis-associated antibodies used were anti-RIP1 (#3493, Cell Signaling Technology), anti-phospho-RIP1 (Ser166, #53286, Cell Signaling Technology), anti-RIP3 (#3493S, Cell Signaling Technology), anti-phospho-RIP3 (Thr231/Ser232, #91702, Cell Signaling Technology), anti-MLKL (#28640S, Cell Signaling Technology), anti-phospho-MLKL (Ser345, #37333, Cell Signaling Technology), and anti-ZBP1 (AG-20B-0010, AdipoGen). The apoptosis-associated antibodies used were anti-cleaved caspase3 (#9664, Cell Signaling Technology) and anti-cleaved caspase8 (Asp387, #8592, Cell Signaling Technology). Other antibodies used were anti-Gr1 (GB11229, Servicebio), anti-CD3 (GB11014, Servicebio), anti-CD45 (GB11066, Servicebio), anti-F4/80 (GB11027, Servicebio), anti-lysozyme (ab108508 Abcam), anti-SDMA (symmetric di- methyl arginine motif, #13222, Cell Signaling Technology), anti-RIP3^479^ (homemade), anti-V5 (#13202, Cell Signaling Technology), anti-HA (#3724, Cell Signaling Technology), and anti-Flag (1001770839, Sigma).

### Histology

Freshly isolated 2-cm small intestine segments were fixed in 4% formalin overnight and then embedded in paraffin, and 5-μm section slides were stained with eosin and hematoxylin. For immunohistochemistry, dewaxed and rehydrated sections were incubated in heat-induced antigen retrieval solution (P0081, Beyotime). The primary antibodies used for staining were anti-cleaved caspase 3, anti-lysozyme, anti-CD3, anti- CD45, anti-Gr1, and anti-F4/80. Biotinylate secondary antibody and DAB substrate were purchased from Boster. For electron microscopy, 3-mm-long samples from the distal, medial and proximal small intestine were excised from *Prmt5*^fl/fl^ and *Prmt5* ^IEC-KO^ mice and fixed in 2.5% glutaraldehyde in phosphate buffer pH 7.4 for 4 h at room temperature. The tissues were embedded in Epon resin after fixation in 1% OsO4.

Then, 70-nm sections were stained with uranyl acetate and lead citrate. Representative pictures were captured using a HITACHI, HT-7800 (JAPAN). An experienced pathologist evaluated cell death on histological sections.

### Cell death assay

For induced necroptosis, cells were seeded in 6-well plates. Cells were pretreated for 3 hours with 20 ng ml^-1^ TNFα (HY-P7090, MedChemExpress) and the pan-caspase inhibitor Z-VAD-FMK (S7023, Selleck, 20 μM) when the cell density was 80%. For necroptosis inhibition, the RIP1 kinase inhibitor necrostatin-1 (Nec-1, HY-15760, MedChemExpress, 5 μM) was used, and the RIP3 kinase inhibitor GSK-872 hydrochloride (HY-101872A, MedChemExpress, 5 μM) was used. For methyltransferase activity inhibition of PRMT5, JNJ-64619178 (HY-101564, MedChemExpress, 0.5 μM), also named onametostat, was used. Pretreated cells were stained with a cell death kit (C10625, Beyotime), and then the ratio of cell death was determined by flow cytometry (006, CytoFLEX, Backman). PI+ cells were calculated.

### TUNEL staining

The cell death of MEFs, L929 cell lines and intestinal tissue sections was determined by TUNEL staining (G3250, Promega) according to the manufacturer’s protocol. In brief, the dewaxed and hydrated slides were incubated in protein K working solution (20 μg ml^-^) for 10 min, and then equilibration buffer (200 mM potassium cacodylate, 25 mM Tris-HCl, 0.2 mM DTT, 0.25 mg ml^-^ BSA, 2.5 mM cobalt chloride) was used for incubation for 5 min. After that, reaction solution (fluorescein-labelled-12-dUTP, rTdT) was used. Link, 1X SSC was used to stop the reaction, then VECTASHIELD® + DAPI (Vector Lab Cat. # H-1200) was used to mount slides.

### Quantitative RT–PCR

Total RNA was extracted with Triol Reagent (Takara), and cDNA was synthesized with Superscript III cDNA-synthesis Kit (Takara). SyBrGreen (Takara) was used to perform RT‒PCR. The β-actin gene was used as a reference gene. The mouse-specific primers were as follows:

Lysozyme P forward, 5’- GCCAAGGTCTAACAATCGTTGTGAGTTG-3’; Lysozyme P reverse, 5’-CAGTCAGCCAGCTTGACACCACG-3’;

Ang4 forward, 5’-GGTTGTGATTCCTCCAACTCTG-3’; Ang4 reverse, 5’-CTGAAGTTTTCTCCATAAGGGCT-3’; Defa-rs2 forward, 5’-ATGAAGAAACTTGTCCTCCTC-3’; Defa-rs2 reverse, 5’-TTATTTTGGATTGCATTTGCA-3’;

Cryptidin1 forward, 5’-TCAAGAGGCTGCAAAGGAAGAGAAC-3’; Cryptidin 1 reverse, 5’-TGGTCTCCATGTTCAGCGACAGC-3’;

TNF forward, 5’-ACCCTGGTATGAGCCCATATAC-3’; TNF reverse, 5’-ACACCCATTCCCTTCACAGAG-3’;

### Depletion of commensal bacteria

Six- to eight-year-old mice were treated with 1 g ampicillin (Boster), 1 g metronidazole (Boster), 500 mg vancomycin (Boster), and 1 g neomycin sulfate (Boster) per liter of drinking water 1 week before tamoxifen injection. Drinking water was refreshed every 2-3 days until the mice died.

### Necrostatin-1 and GSK-872 administration

For pregnant mice, 50 mg kg^-^per body weight necrostatin-1 was applied by intraperitoneal injection every day from offspring E19.5 to E15.5. Six- to eight-week-old mice received 70 mg kg-per body weight necrostatin-1 with tamoxifen by intraperitoneal injection every day until death. GSK-872 (10 mg kg^-^ per body weight) was applied with tamoxifen by intraperitoneal injection every day. Both reagents were dissolved in 5% DMSO+ 40% PEG300 +5% Tween 80 + 50% ddH2O.

### Global mapping of PRMT substrates

The IEC samples were isolated from 4-week-old *Prmt5*^fl/fl^ (n=6) and *Prmt5*^IEC-KO^ (n=6) mice by sequentially incubating intestinal tissue in 1 mM DTT and 1.5 mM EDTA solutions as described previously(Wei et al., 2020). The prepared tissue was lysed in modified SDT buffer (0.1 M Tris-HCL, pH 7.6, 0.1 M DTT, 1% SDS, 1% SDC), which were used to map the PTMScan asymmetric di- or symmetrical di-methyl arginine motif according to a protocol described previously (Li et al., 2021). Briefly, the lysate was alkylated in UA solution (8 M urea, 100 mM Tris-HCl pH 8.5) after genomic DNA was sheared and ultrafiltrated. Then, the samples were digested by Lys-C and trypsin, acidified (pH

2.0) by formic acid, loaded onto SepPak tC18 cartridges, desalted and eluted in 70% acetonitrile, lyophilized and stored before use. For methylome analysis, 15 mg peptides were off-line fractionated by basic reversed-phase, and 60 fractions were concatenated to 10. One microgram of peptides was used for each run after all fractions were concentrated and lyophilized. PTMScan asymmetric di- or symmetrical di-methyl arginine motif immunoaffinity beads (CST, 13563, and 13474) were used to enrich associated peptides. All LC‒MS samples were performed with an EASYnLC 1200UHPLC system (Thermo Fisher Scientific) and analysed with Xcalibur software (Thermo Fisher Scientific, v4.0). Raw data processing in Proteome Discoverer (Thermo Fisher Scientific, v2.2) and di-methylation sites were extracted by the IceLogo web server according to 5 upstream amino acids and 11 downstream amino acids of the arginine methylation site identified in total(Colaert et al., 2009).

### Coimmunoprecipitation

Plasmids encoding RIP3, RIP1, MLKL, full-length ZBP1, and RHIM-mutation ZBP1 were kind gifts from Professor Jiahuai Han and Professor Wei Mo at Xiamen University. The RIP3 (RIP3-R479K) fusion protein was constructed from the corresponding template. Plasmids encoding PRMT5 and the mutant PRMT5 (PRMT5-E444Q) were constructed in-house. All constructs were cloned and inserted into PCDH vectors with V5, Flag and HA tags. The sequences of all constructs were confirmed by DNA sequencing. HEK293-T and L929 cells were transfected with the indicated plasmids by PEI (homemade) or Lip 8000 (C0533, Beyotime). Cells were harvested at 48 hours after transfection. Immunoprecipitation assays were performed as described previously(Zhang et al., 2002). Briefly, cells were lysed with lysis buffer (P0013, Beyotime) on ice for at least 4 hours, and the supernatant was collected by centrifugation. Equal amounts of protein from samples were reserved for analysing protein expression. The remaining protein was separately incubated with anti-V5 magnetic beads (P2141, Beyotime), anti-HA magnetic beads (B26202, Selleck), and anti-Flag magnetic beads (26201, Selleck) at 4°C overnight, and anti-protein A/G magnetic beads were used as a negative control. Second, the beads were collected by magnetic stand (Selleck), washed with PBST 3 times, and incubated with 1x protein loading buffer (Beyotime) to release immunoprecipitates. Total protein and immunoprecipitates were resolved by SDS‒PAGE for immunoblotting analyses.

### Confocal microscopy

Cell immunostaining was described previously(Chen et al., 2022a). Briefly, freshly prepared 4% formaldehyde was used to fix cells for 15 min at room temperature, and then 0.5% Triton-100 was used for permeabilization. After blocking with 5% BSA, the samples were incubated with primary antibody at 4°C overnight and then stained with secondary antibody for 1 hour at room temperature. Finally, the samples were mounted with anti-fluorescence fade reagent (P36934, Invitrogen). The negative control included samples stained without primary antibodies and samples with no signals of an interesting protein.

The images were captured using a laser scanning confocal microscope (LSM 780, Zeiss).

### STORM imaging and analysis

An N-STORM microscope (Ti-E, Nikon) was used to perform the STORM imaging, according to previously described methods(Chen et al., 2022a). In brief, the N-STORM system included an Agilent MLC-400B laser launch with 4 color lasers (a 674 nm red diode laser, a 561 nm green solid-state laser, a 488 nm blue solid-state laser, and a 405 nm violet diode laser) and a 100 X oil immersion objective. The emission fluorescence was detected by a back-illuminated EMCCD camera (iXon DU897, Andor) after separation using appropriate filters (FF02-520/28-25, FF01-586/20-25x3.5 and FF01- 692/40-25; Semrock). Samples were labelled with interesting antibodies and then immersed in imaging buffer (50 mM pH 8.0). Tris, 10 mM NaCl, 0.5 mg ml^−1^ glucose oxidase, 40 μg ml^−1^ catalase, 10% glucose and 143 mM β-mercaptoethanol). For signal- color STORM imaging, the CF 647 channel was recorded by exposure to a 647 nm laser at a power density of 2 kW cm^−2^. For two-color STORM imaging, CF 647 and CF 568 channels were recorded in turn and then reconstructed by the N-STORM module in NIS-Elements AR software. Ilastik, a machine machine-learning-based segmentation toolkit, was used to analyse the cluster, as described previously(Chen et al., 2022a). Briefly, based on the feature description of high-quality STROM images, the classification of effective signals was determined. Each effective cluster was strictly identified as a separate object. Through evaluating the segmentation outcomes, analysis parameters were finally determined and used for all images. Fiji/ImageJ was used to process the output binary images by extracting the area of effective clusters and Feret parameters. To distinguish subunits of RIP1/RIP3 clusters in the same complex, the property of coclusters in single-colour or overlap between two-colour images was determined by the distance of the centroid in each subunit and their area.

### Modelling and MD details

The coordinates of RIP1_Mouse (accession number: Q60855) and RIP3_Mouse (accession number: Q9QZL0) proteins were modelled using the AlphaFold prediction method (Jumper et al., 2021). All the heteroatoms of nonprotein parts were removed, and missing hydrogen atoms were added with the expected protonation states of residues (Accelrys, 2011). The three protein structures (RIP1 and RIP3) were geometrically optimized using the conjugate gradient (CG) method and further refined by 100-ns explicit solvent molecular dynamics (MD) simulations using GROMACS 2018.8 (Abraham et al., 2015) and the Charmm36m force field (Huang et al., 2017).

Details of the MD simulation was published previously (Li et al., 2019b; Yang et al., 2022; Zang et al., 2022). In brief, the transferable intermolecular potential with three points (TIP3P) water model (Berendsen et al., 1981) was employed to describe the solvent. The temperature of each system was kept at 300 K using the velocity rescaling coupling method (Bussi et al., 2007). The protein and nonprotein groups were separately coupled to an external heat bath with a 0.1 *ps* relaxation time. The pressure was kept at 1.0 *bar* with a coupling constant of 2.0 *ps* using the Parrinello-Rahman method (Li et al., 1992). Constraints were applied for hydrogen bond lengths using the SETTLE algorithm (Miyamoto and Kollman, 1992) for solvent and the LINCS algorithm (Hess, 2008) for the proteins, which allows a 2.0 *fs* integration time step using the Verlet integrator. Electrostatic interactions were calculated using the particle‒mesh Ewald method with a real space cut-off of 12.0 Å. The van der Waals interactions were treated with a cut-off of 12.0 Å. Cl^−^ was added to neutralize the systems, and then 150 mM NaCl was added to simulate physiological conditions. Periodic boundary conditions were applied in all simulations.

The RIP1-RIP3 heterooctomer was generated by four RIP1 peptides (seq: 520-535) and four RIP3 peptides (seq: 409-486) in accordance with the reported structure of RIP1/RIP3 oligomers (Chen et al., 2022b). The optimal docked complex was selected on the basis of energy and the size of the cluster and optimized using the conjugated gradient (CG) algorithm, with a convergence criterion of 0.01 kcal·mol^−1^·Å^−1^ (Yang et al., 2023; Yang et al., 2022). Residue R479 of RIP3_Mouse was mutated to lysine (RIP3^R479K^) by using Discovery Studio (Accelrys, 2011). Each energy-minimized docked complex was further refined by 100-ns MD simulations (Yang et al., 2023).

The binding energies (*ΔGbind*) of RIP3 (RIP3^R479K^) with RIP1 were estimated based on 500 snapshots evenly extracted from 50∼100 ns MD trajectories (Yang et al., 2023; Yang et al., 2021). The exterior (solvent) and interior dielectric constants were set to 80.0 and 1.0, respectively (Wang et al., 2019). The data analysis was performed using the tools included in the GROMACS 2018.8 software package and our in-house codes. Structural plotting and visualization were accomplished by Discovery studio client (Accelrys, 2011) and the Visual Molecular Dynamics (VMD) program (Humphrey et al., 1996).

## Funding

This study was funded by the National Natural Science Foundation of China (No. 82073202), the Fund for Scientific and Technological Innovation Team of Shaanxi Innovation Capability Support Plan (2023-CX-TD-67), and the State Key Laboratory of Cancer Biology Project (CBSKL2022ZZ15).

## Acknowledgement

We would like to thank professor Jiahuai Han and Wei Mo of Xiamen University for helpful providing the *mlkl* and *Zbp1* knockout mice, and discussions on topics related to this work. We thanks for professor Feng Shao of NIBS, Beijing for providing the conditional *Gsdmd* knockout mice and professor Xin Lin of Tsinghua University providing the *Tnfr1* knockout mice. We are grateful to Dr. Peng Lin of Fourth Military Medical University for his help with protein structure analysis discussion.

---

**Figure S1.**
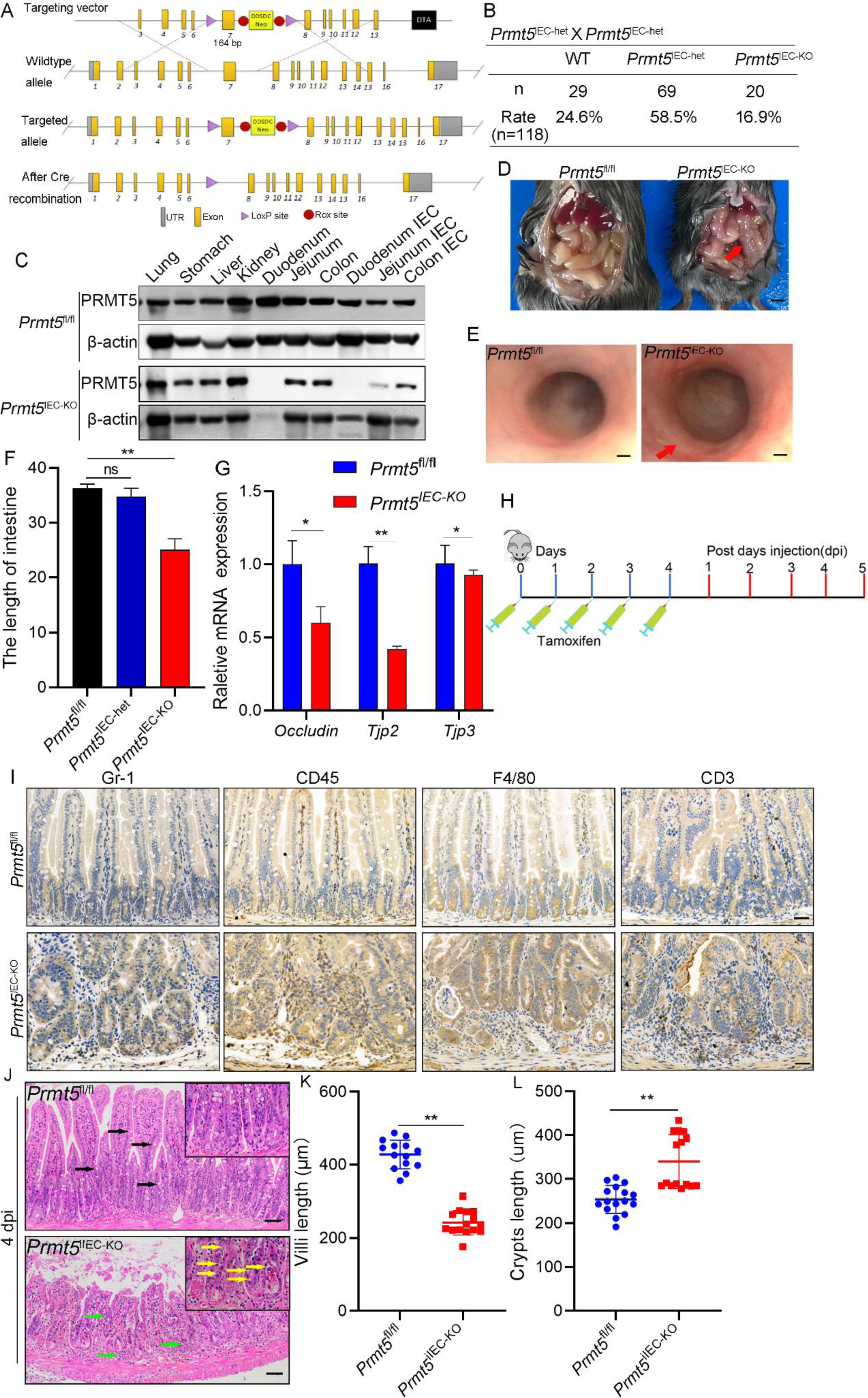
Characterization of *Prmt5*^IEC-KO^ mice. A, Schematic representation of the targeting vector along with wildtype and targeted allele in *Prmt5* ^fl/fl^ mice; B, Birth rates of all genotypes of offspring from selected breeding couples (*Prmt5*^IEC-^ het _X *Prmt5*_IEC-het_);_ C, Immunoblotting of PRMT5 expression in various tissues of *Prmt5*^IEC-KO^ mice and control *Prmt5* ^fl/fl^ mice; D, Anatomy images of intestine of *Prmt5*^IEC-KO^ mice and control *Prmt5* ^fl/fl^ mice at 3weeks; Red arrow indicted signs of watery diarrhea; E, Endoscopic images of *Prmt5*^IEC-KO^ mice and control *Prmt5* ^fl/fl^ mice at 3weeks; Red arrow showed irregular surface and lesions; F, Statistical analysis for the length of intestine in *Prmt5*^IEC-KO^ and *Prmt5*^IEC-het^ mice compared with *Prmt5* ^fl/fl^ mice, n=3; *P<0.05, **P<0.01; G, Quantitative real-time PCR (qPCR) analysis of intestinal epithelium cells tight junctions associated genes *Occludin*, *Tjp2* and *Tjp3* on RNA extracted from small intestine mucosa of *Prmt5*^IEC-KO^ (n=3) and control *Prmt5*^fl/fl^ (n=3) mice; *P<0.05, **P<0.01; H, Schematic representation of tamoxifen injection scheme at 6-8 weeks mice; I, Lymphocyte IHC signals in small intestine of *Prmt5*^IEC-KO^ (n=3) and *Prmt5*^fl/fl^ (n=3) mice; Scale bars, 100 µm; J, Hematoxylin and eosin staining (HE) of small intestine sections from *Prmt5*^iIEC-KO^and *Prmt5*^fl/fl^ mice at 4dpi; Black arrows, goblet cells; Green arrows, infiltrating leukocytes; Yellow arrows, dead cells; Scale bars, 100 µm; K, Statistical analysis for lengths of small intestine villi; *P<0.05, **P<0.01. L, Statistical analysis for lengths of small intestine crypts; *P<0.05, **P<0.01.

**Figure S2.**
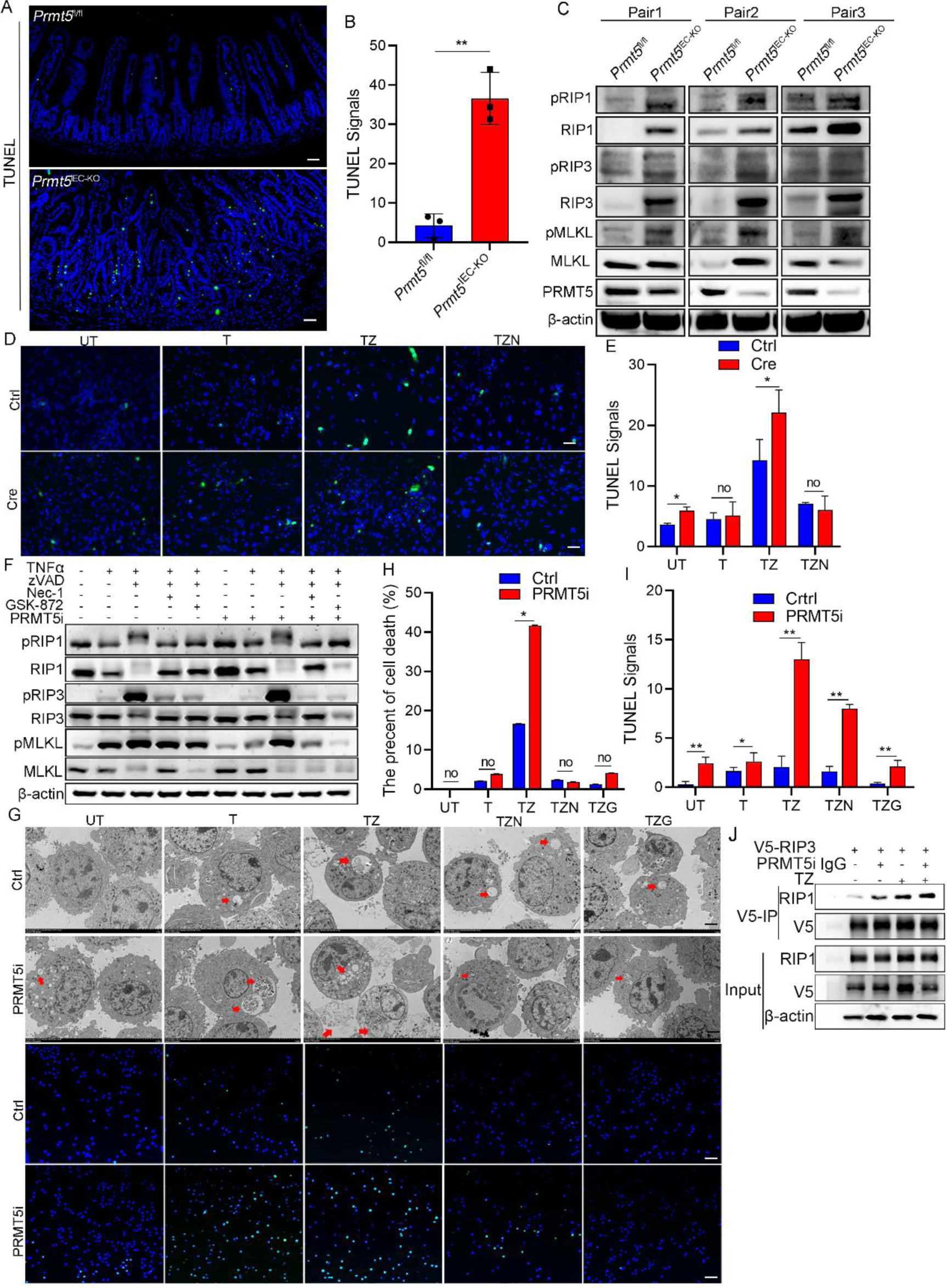
Epithelial PRMT5 depression triggers necroptosis A, TUNEL assay (green) in small intestine of *Prmt5*^IEC-KO^ and control *Prmt5*^fl/fl^ mice; B, Statistical analysis for TUNEL signals in small intestine of *Prmt5*^IEC-KO^ and control *Prmt5*^fl/fl^ mice; C, Immunoblotting of necroptosis signaling in IECs of *Prmt5*^IEC-KO^ and control *Prmt5*^fl/fl^ mice, n=3; D, TUNEL assay (green) in MEFs; E, Statistical analysis for TUNEL signals in MEFs; *P<0.05, **P<0.01. F, Immunoblotting of necroptosis signaling in L929 cells; G, Electron microscopy images of necrotic-like IECs and TUNEL assay (green) in L929 cells. UT: Untreated; T: TNFα; TZ: TNFα+zVAD-FMK; TZN: TNFα+zVAD-FMK+Nec-1; TZG: TNFα+zVAD-FMK+GSK-872. Red arrows: necrotic-like IECs. TEM, transmission electron microscopy; H, Statistical analysis for necrotic-like IECs in L929 cells; *P<0.05, **P<0.01. I, Statistical analysis for TUNEL signals in L929 cells; *P<0.05, **P<0.01. J, V5-RIP3 were expressed in RIP3-KO L929 cells treated with or without PRMT5i and TZ, immunoprecipitated with anti-V5 beads, followed by immunoblotting with anti-RIP1 antibody;

**Figure S3.**
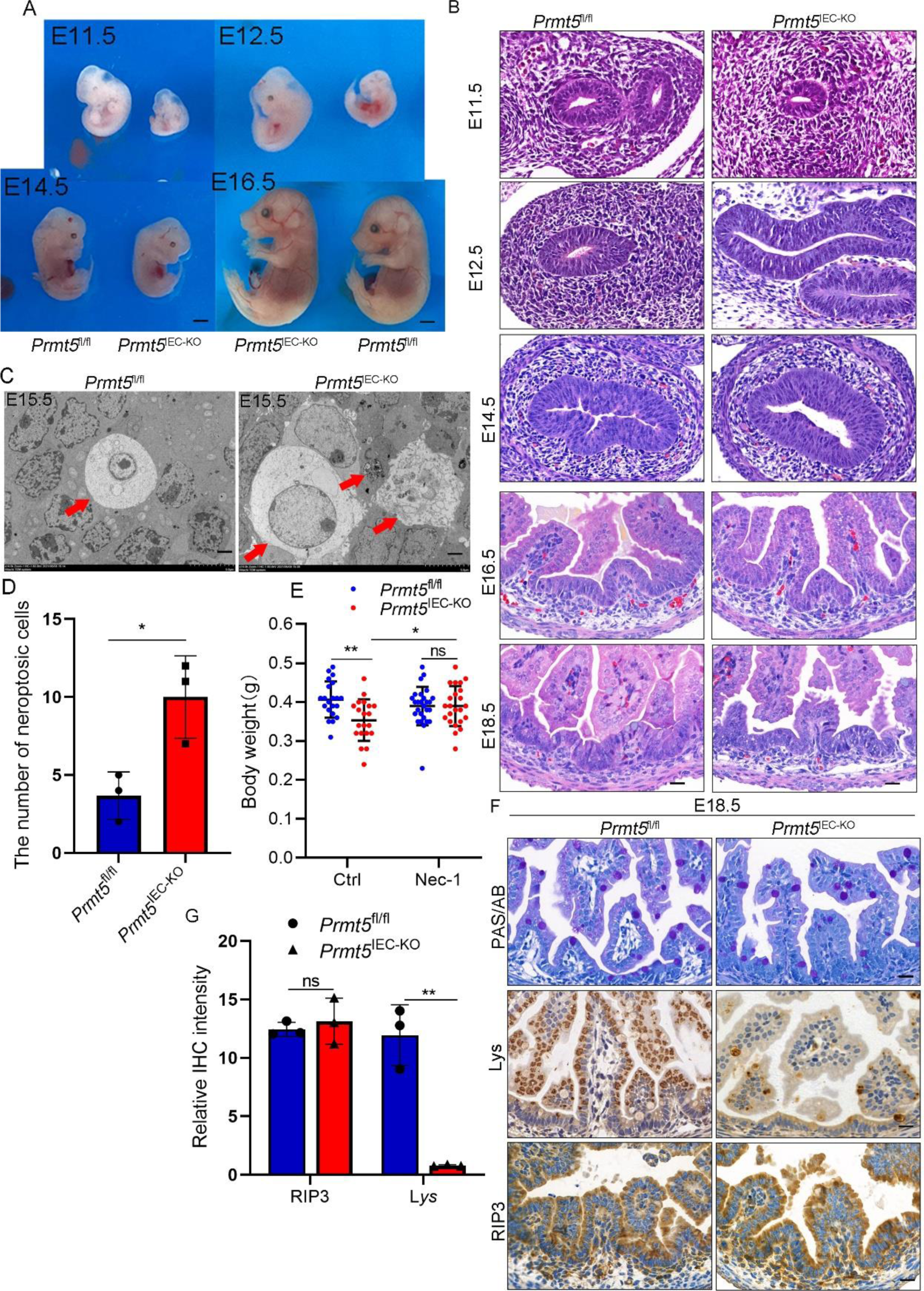
*Prmt5* deficiency mice exhibited normal intestinal development at all embryonic stages, but high necroptotic level in intestine A, Macroscopic features of *Prmt5*^IEC-KO^and *Prmt5*^fl/fl^ mice at E11.5, E12.5, E14.5, E16.5; Scale bars, 100 µm; B, Haematoxylin and eosin staining (HE) of small intestine sections from *Prmt5*^IEC-KO^ and *Prmt5*^fl/fl^ mice at E11.5, E12.5, E14.5, E16.5, E18.5; Scale bars, 100 µm; C, Electron microscopy images of necrotic-like IECs in *Prmt5*^IEC-KO^ (n=3) mice at E15.5; Red arrows: necrotic-like IECs. TEM, transmission electron microscopy; D, Statistical analysis for necrotic-like IECs in *Prmt5*^IEC-KO^ mice; *P<0.05, **P<0.01. E, Body weight of *Prmt5*^IEC-KO^ and *Prmt5*^fl/fl^ mice at E15.5; Nec-1: Necrostatin-1; F, Periodic acid–Schiff/Alcian blue (PAS/AB; goblet cells) staining, lysozyme (Lys) (Paneth cells) and RIP3 immunohistochemical staining in the small intestine. Three individual experiments were repeated; Scale bars, 100 µm; G, Statistical analysis for Lys and RIP3 IHC signals; *P<0.05, **P<0.01.

**Figure S4.**
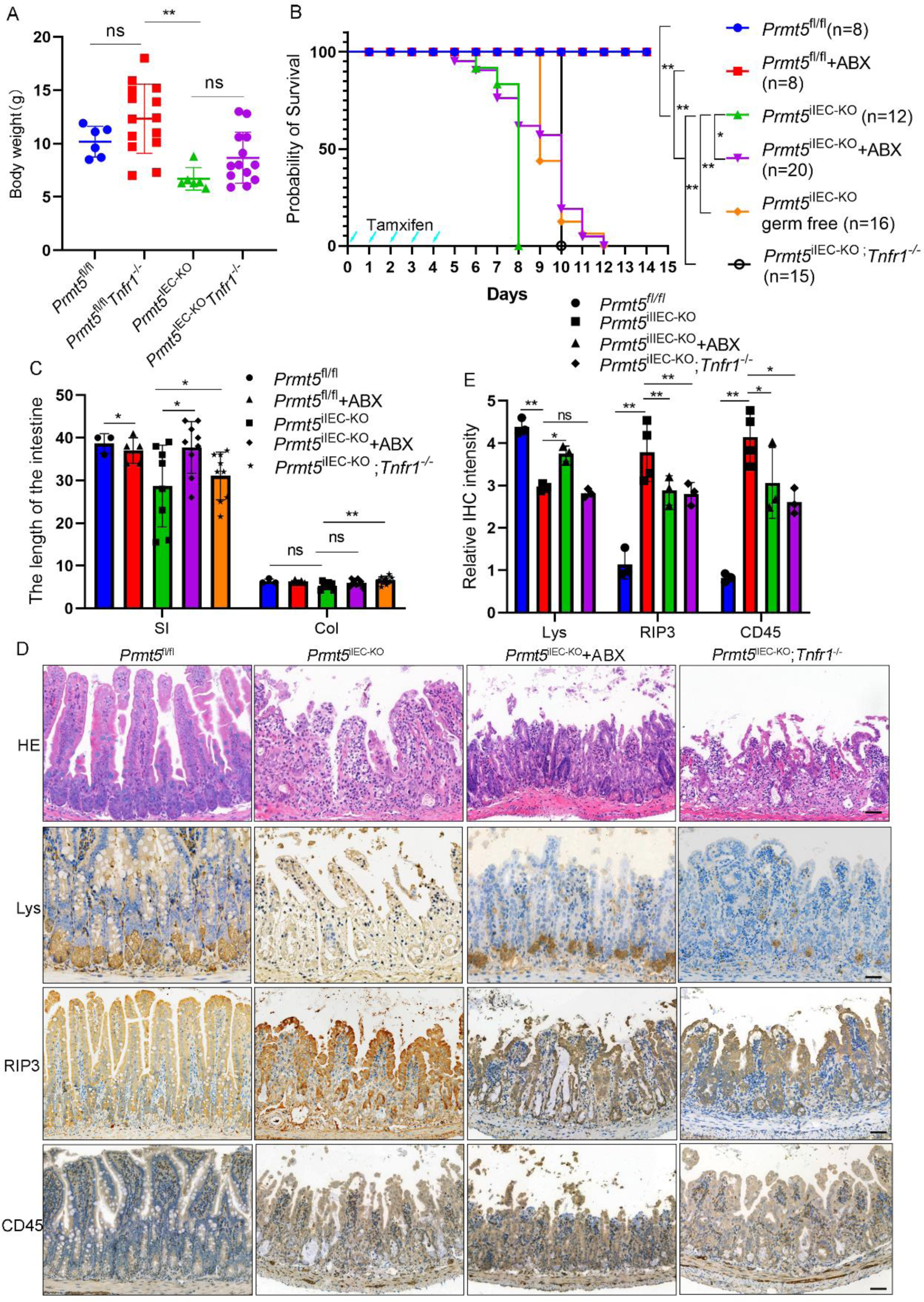
*Prmt5*^iIEC-KO^ phenotype in conditions of antibiotic treatment, TNFR1 deficiency or germ-free mice. A. Body weight of *Prmt5*^IEC-KO^, *Tnfr1*^-/-^, *Prmt5*^IEC-KO^ *Tnfr1*^-/-^ DKO and control *Prmt5* ^fl/fl^ m; *P<0.05, **P<0.01 B, Survival rates were compared among different groups: *Prmt5*^iIEC-KO^ *Tnfr1*^-/-^, *Prmt5*^iIEC-KO^ mice treated with broad-spectrum antibiotics (ABX), *Prmt5*^iIEC-KO^ germ-free and *Prmt5* ^fl/fl^ mice; *P<0.05, **P<0.01 C, Statistical analysis for the gut length in different groups of mice. SI (small intestine), Col (colon); *P<0.05, **P<0.01 D, Representative images of small intestine sections from mice with the indicated genotypes stained with HE or immunostained for Lys, RIP3and CD45; Scale bar, 100 μm E, Statistical analysis for IHC signals of Lys, RIP3 and CD45; *P<0.05, **P<0.01

**Figure S5.**
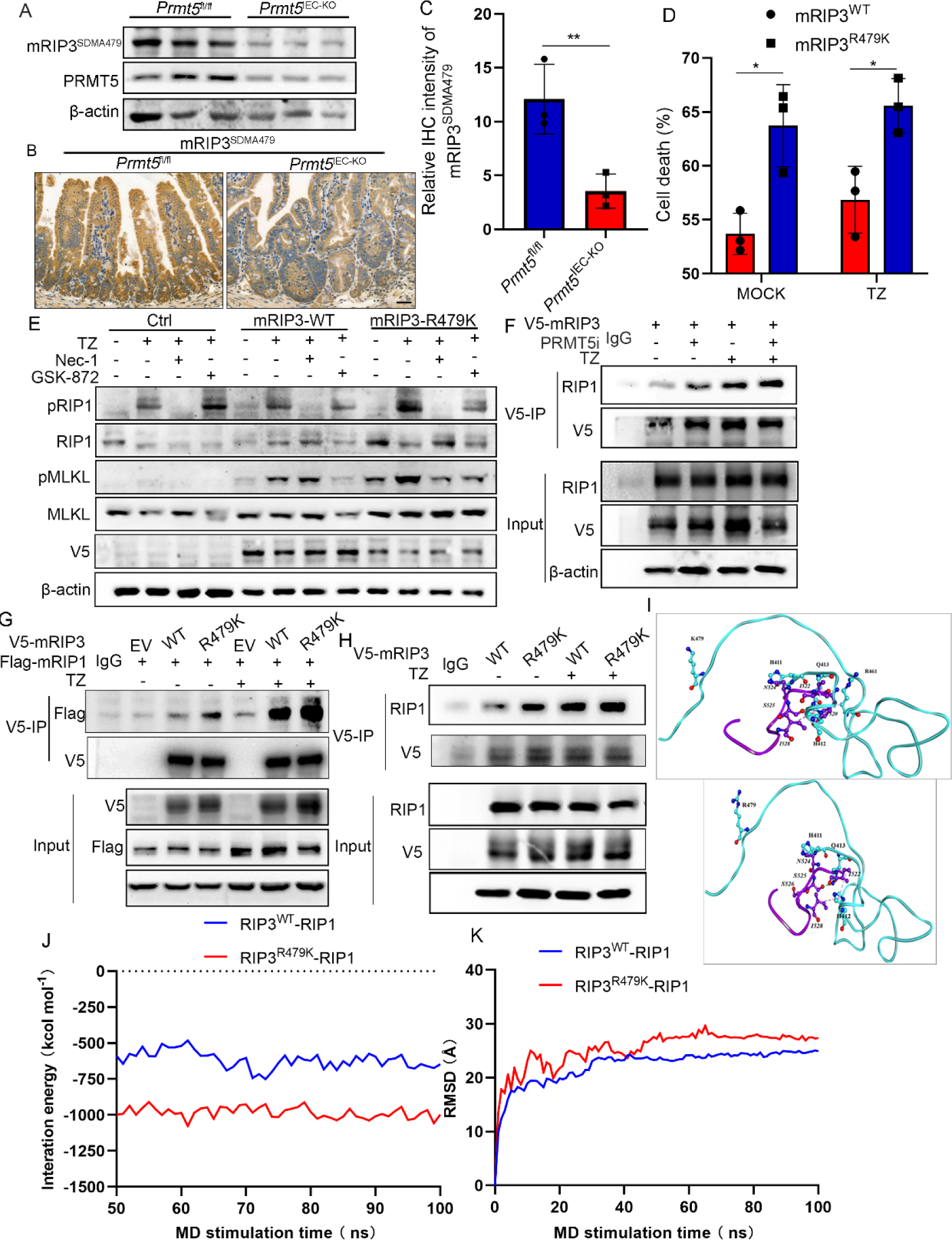
Arginine methylation of RIP3 on R479 suppresses necrosome A, Immunoblotting of RIP3^SDMA479^in IECs derived from *Prmt5*^IEC-KO^ (n=3) and control *Prmt5*^fl/fl^ (n=3) mice; B, RIP3^SDMA479^ IHC signals in small intestine of *Prmt5*^IEC-KO^ (n=3) and *Prmt5*^fl/fl^ (n=3) mice; Scale bars, 100 μm; C, Statistical analysis for RIP3^SDMA479^ IHC signals in *Prmt5*^IEC-KO^ mice. *P<0.05, **P<0.01; D, Cell viability was measured on the basis of quantitation of the flow cytometry; *P<0.05, **P<0.01; E, Immunoblotting of phosphorylation level of RIP1 and MLKL in L929 cells which overexpressed RIP3-WT or 479 mutate and induced necroptosis signaling by TZ combined with Nec-1 or GSK-872; F, V5-RIP3 WT were expressed in RIP3-KO L929 cells treated with or without PRMT5i and TZ (20 ng^-ml^ TNFα, 20 μM zVAD), then immunoprecipitated with anti- V5 beads, followed by immunoblotting with anti-RIP1 antibody; G, Both Flag-RIP1 and V5-RIP3 WT or mutant R479K overexpressed in 293T cells with or without necroptosis induction by TZ (T: 20 ng^-ml^ TNFα; Z: 20 μM zVAD), then immunoprecipitated with anti-V5 beads, followed by immunoblotting with anti- V5 and anti-Flag antibody; H, V5-RIP3 WT or mutant R479K overexpressed in RIP3-KO L929 cells with or without necroptosis induction by TZ (T: 20 ng^-ml^ TNFα; Z: 20 μM zVAD), then immunoprecipitated with anti-V5 beads, followed by immunoblotting with anti-RIP1 antibody; I, Propeller structure and key residues within the binding interface of RIP3-RIP1 and RIP3^R479K^-RIP1 system. The key residues are represented by ball and stick models, and the important H-bonding (or electrostatic) interactions are labeled in the dashed green lines. The H atoms are omitted for clarity; J, Binding energies (*ΔGbind*) of the four systems which were estimated on the basis of 500 snapshots evenly extracted from 50 ∼ 100 ns MD trajectories; K, Backbone-atom root-mean-square deviations (RMSD) of the four systems over the 100-ns molecular dynamics (MD) simulations.

**Figure S6.**
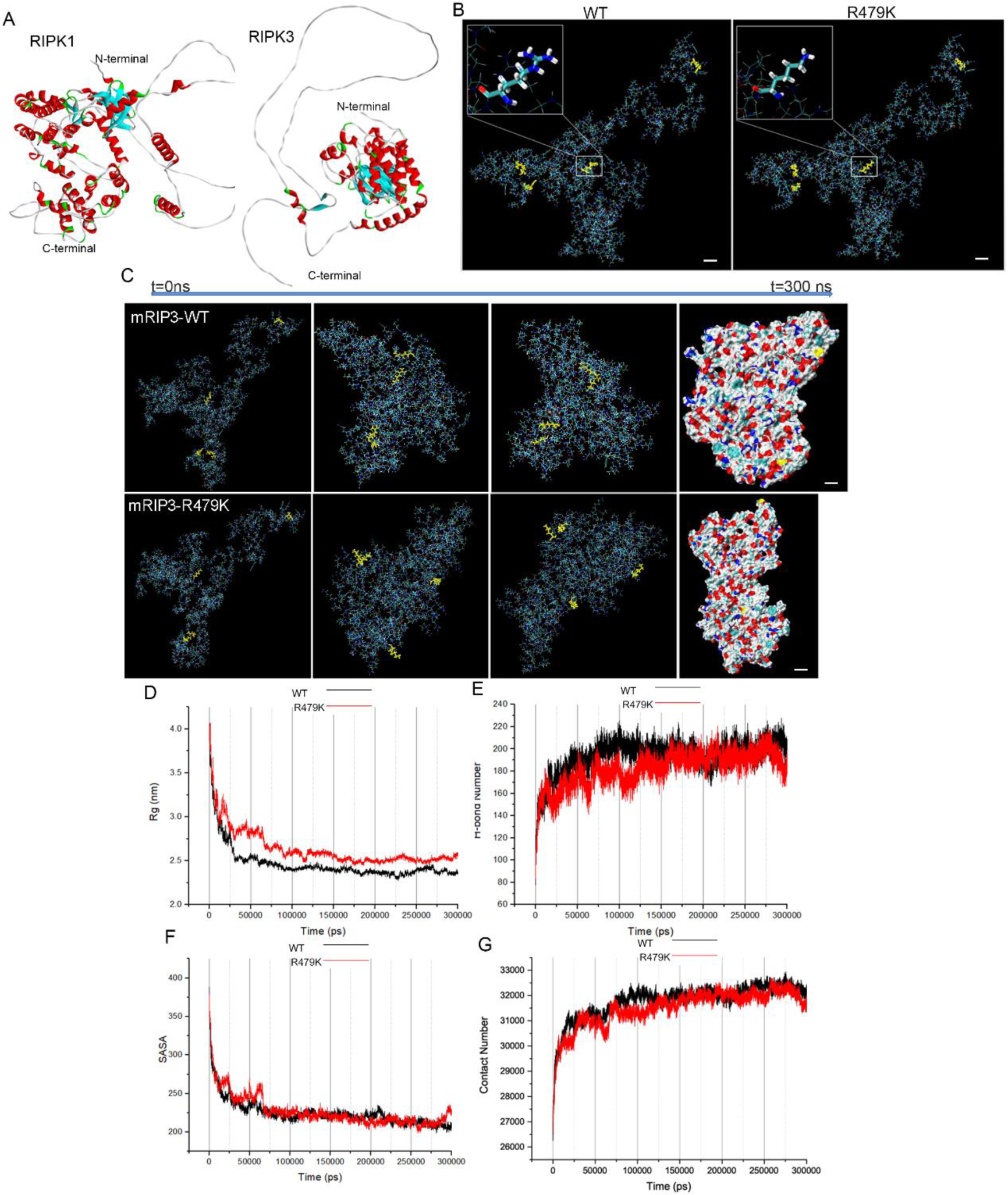
RIP3-RIP1 modeling and MD details A, Modeling structures of the RIP1 and RIP3 proteins after the 100-ns molecular dynamics (MD) simulations. The initial models are conducted by the AlphaFold prediction method. The colors of the ribbons distinguish between helices (red), β- sheets (cyan), hydrogen-bonded turns (green), and random coils (white); B, Propeller structure and key residues within the binding interface of RIP3-RIP1 and RIP3^R479K^-RIP1 system; C, The time evolutional snapshots of WT and R479K mutant conformations. The mutation sites are marked in yellow; D, The time evolution of the radius of gyration (Rg) of WT and R479K mutant; E, The time evolution of hydrogen bond (H-bond) probability of WT and R479K mutant; F, The time evolution of solvent accessible surface area (SASA) of WT and R479K mutant; G, The time evolution of Contact number of WT and R479K mutant.

**Figure S7.**
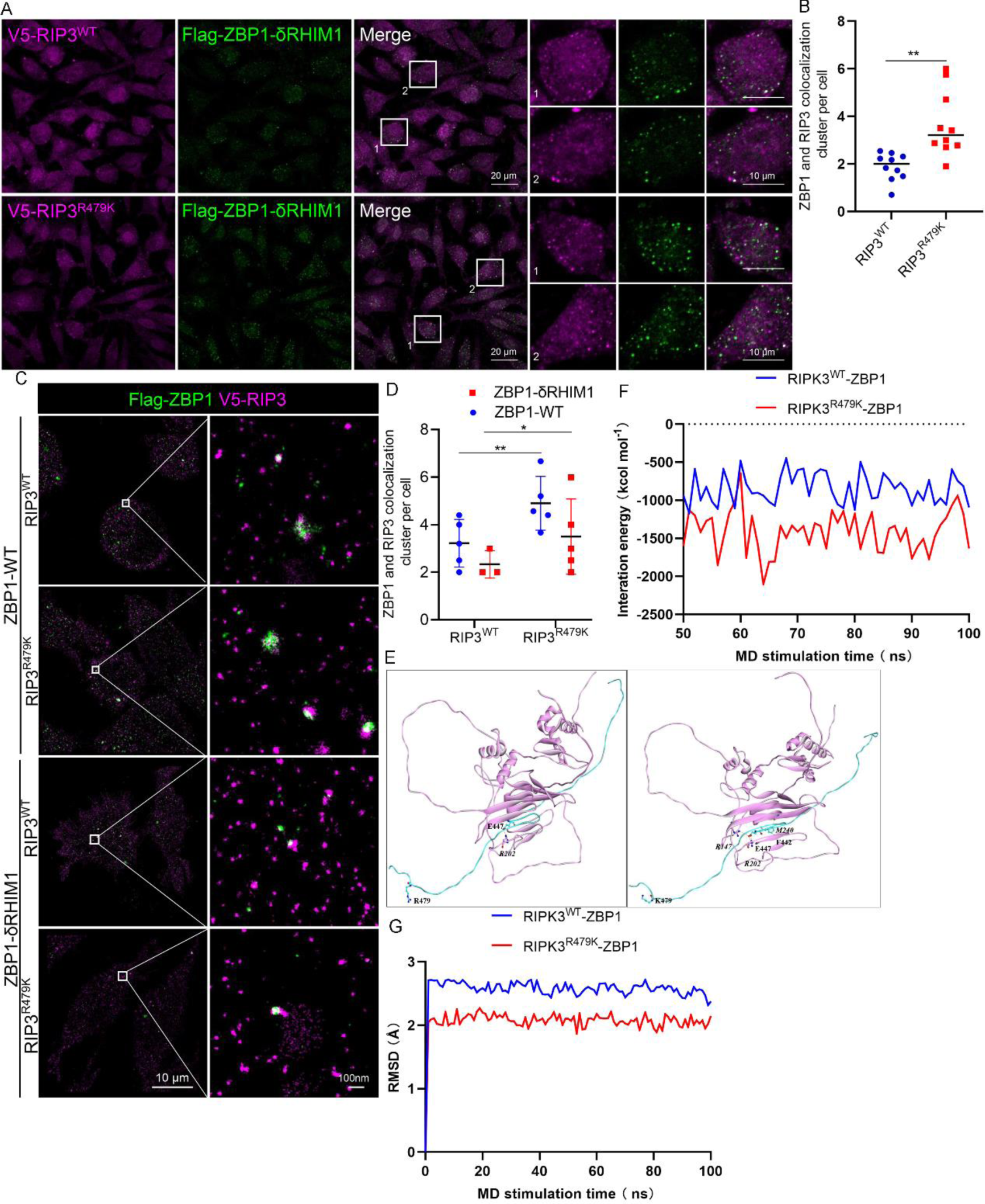
Arginine methylation of RIP3 on R479 suppresses combination between RIP3 and ZBP1 A, Two-colour confocal images of RIP3 WT or mutant R479K and ZBP1 mutant ΔRHIM1 in cells, the white box magnified images highlighting the RIP3 and ZBP1 colocalization; B, Statistical analysis for RIP3 and ZBP1 colocalization cluster per cell. *P<0.05, **P<0.01; C, Two-colour STORM images showing RIP3 WT or mutant R479K and ZBP1 WT or mutant ΔRHIM1 in L929 cells, the white box magnified images highlighting the RIP3 and ZBP1 colocalization; D, Statistical analysis for RIP3 and ZBP1 colocalization cluster per cell. *P<0.05, **P<0.01; E, Propeller structure and key residues within the binding interface of (A) RIP3-ZBP1 and RIP3^R479K^-ZBP1 system. The key residues are represented by ball and stick models, and the important electrostatic interactions are labeled in the dashed orange lines. The H atoms are omitted for clarity; F, Binding energies (*ΔGbind*) of the four systems which were estimated on the basis of 500 snapshots evenly extracted from 50 ∼ 100 ns MD trajectories; G, Backbone-atom root-mean-square deviations (RMSD) of the four systems over the 100-ns molecular dynamics (MD) simulations.

**Figure S8.**
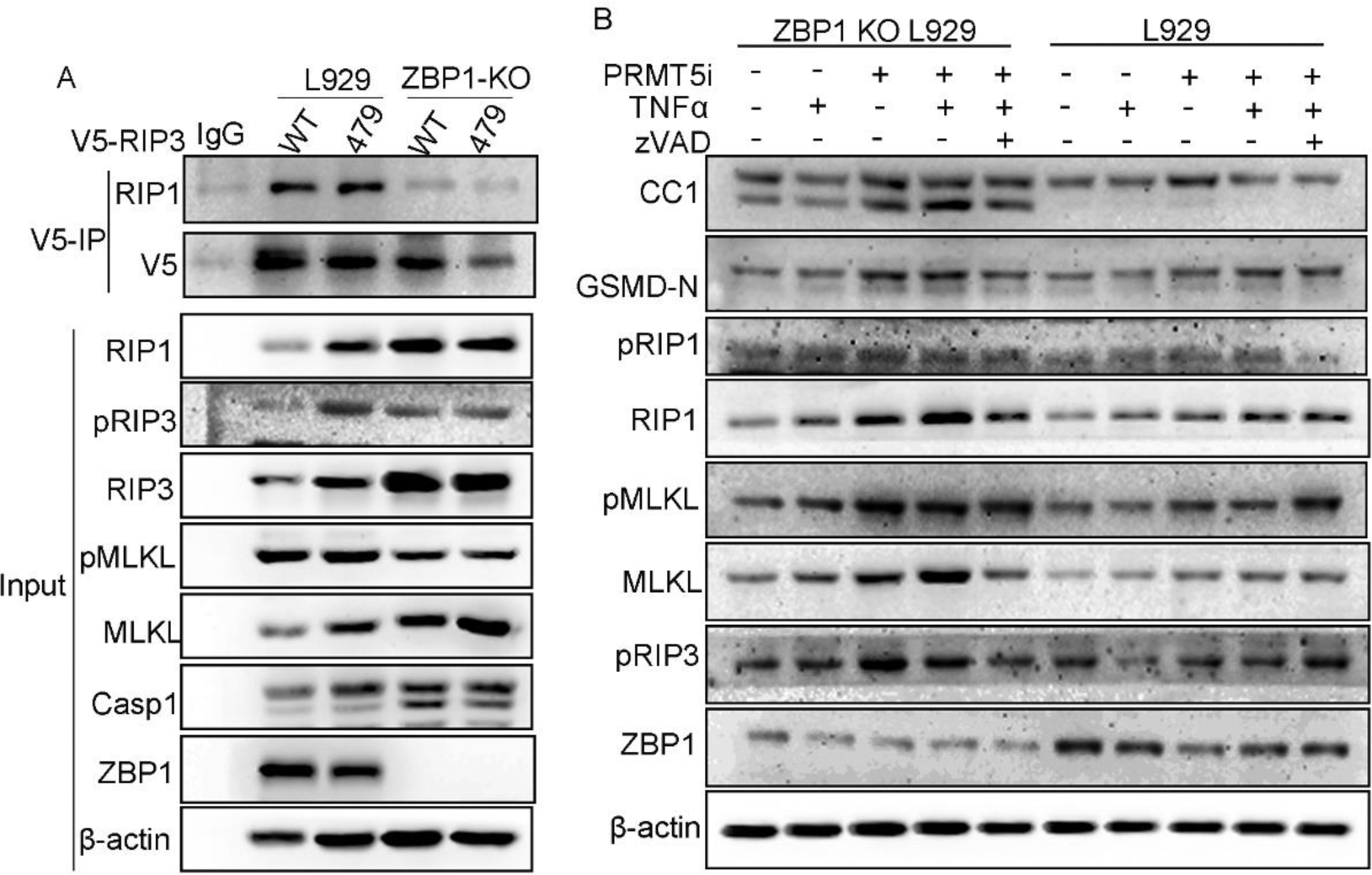
A, V5-RIP3 WT or mutant R479K were overexpressed in L929 or ZBP1-KO L929 cells. Immunoprecipitated for the indicated intrinsic RIP1 and immunoblotting for cell death signaling in cell lysate; B, Immunoblotting of necroptosis and pyroptosis signaling molecules in L929 or ZBP1- KO L929 cells treated as indicated (DMSO, 0.5 μM PRMT5i, 20 ng^-ml^ TNFα, 20 μM zVAD).

**Figure S9.**
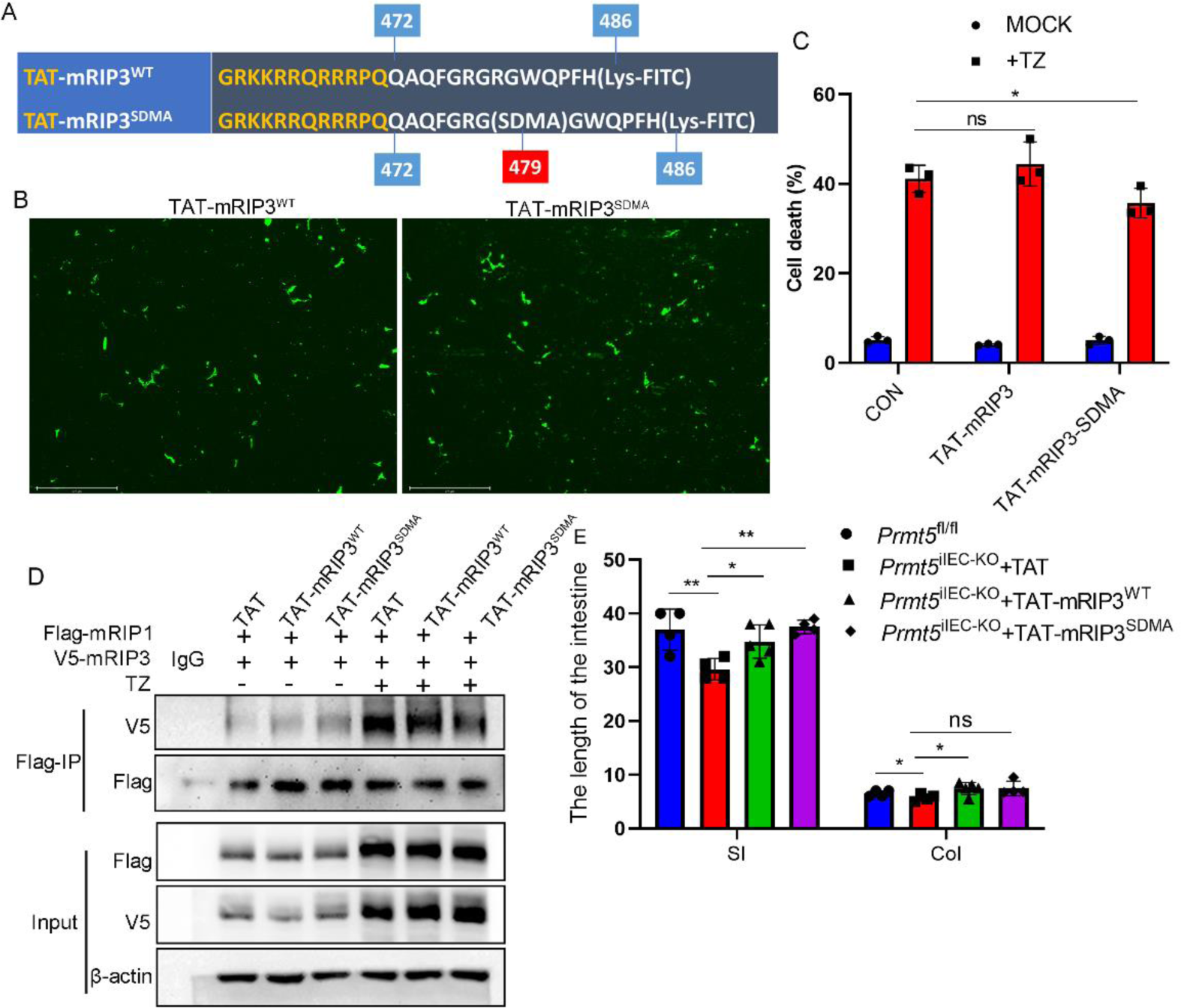
Peptide modified with arginine methylation of RIP3 on R479 suppresses necroptosis mediated by RIP3 A, The amino acids information of TAT-RIP3^WT^ and TAT-RIP3^SDMA^; B, The efficiency of peptides transferred into the targeted cell; C, The precent of cell death after inducing necroptosis by TZ in L929 cells treated with different peptides; D, Both V5-RIP3 and Flag-RIP1 were overexpressed in 293T cells treated with TAT, TAT-mRIP3^WT^ or TAT-mRIP3^SDMA^ peptides with or without necroptosis induced by TZ. Immunoprecipitated with anti-Flag beads. E, Statistical analysis for the gut length in different strains of mice. SI (small intestine), Col (colon); *P<0.05, **P<0.01

## Notes

### Competing Interest Statement

The authors have declared no competing interest.

